# Diverse modes of synaptic signaling, regulation, and plasticity distinguish two classes of *C. elegans* glutamatergic neurons

**DOI:** 10.1101/175836

**Authors:** Donovan Ventimiglia, Cornelia I. Bargmann

## Abstract

Synaptic vesicle release properties vary between neuronal cell types, but in most cases the molecular basis of this heterogeneity is unknown. Here, we compare *in vivo* synaptic properties of two neuronal classes in the *C. elegans* central nervous system, using VGLUT-pHluorin to monitor synaptic vesicle exocytosis and retrieval in intact animals. We show that the glutamatergic sensory neurons AWC^ON^ and ASH have distinct synaptic dynamics associated with tonic and phasic synaptic properties, respectively. Exocytosis in ASH and AWC^ON^ is differentially affected by SNARE-complex regulators that are present in both neurons: phasic ASH release is strongly dependent on UNC-13, whereas tonic AWC^ON^ release relies upon UNC-18 and on the protein kinase C homolog PKC-1. Exocytosis and retrieval each have two timescales in AWC^ON^ but one major timescale in ASH. Strong stimuli that elicit high calcium levels also increase exocytosis and retrieval rates in AWC^ON^, generating distinct tonic and evoked synaptic modes. These results highlight the differential deployment of shared presynaptic proteins in neuronal cell type-specific functions.

## Introduction

Neurotransmitter release is a highly regulated process that varies at different synapses, and at the same synapse over time (Atwood and Karunanithi, 2002). Although presynaptic diversity is widely observed, it is challenging to define its role in intact circuits under physiological patterns of activity (Regehr, 2012); more often, a synapse is examined *ex vivo* at calcium concentrations or temperatures that alter its properties. Studies in the zebrafish retina represent one example in which synapses of two distinct neuronal classes, on- and off-bipolar cells, have been compared *in vivo* in intact animals, leading to insights into their similarities and differences (Odermatt et al., 2012). Here, we extend this approach to the central nervous system of the nematode worm *Caenorhabditis elegans,* and relate diversity in synaptic properties to requirements for specific synaptic proteins in individual neurons.

The well-studied neural circuitry of *C. elegans,* which includes 302 neurons and about 9000 synaptic connections, presents an opportunity to study presynaptic diversity in a well-defined context (Varshney et al., 2011; White et al., 1986). Most synaptic proteins are conserved between *C. elegans* and other animals; indeed, behavioral genetics in *C. elegans* led to the initial identification of the SNARE (soluble N-ethylmaleimide–sensitive factor attachment receptor) regulatory proteins *unc-13* and *unc-18* (Brenner, 1974; Gengyo-Ando et al., 1993; Maruyama and Brenner, 1991). However, the study of synaptic transmission in *C. elegans* has been largely limited to the neuromuscular junction (NMJ) due to the challenges of electrophysiology in this small animal (Richmond and Jorgensen, 1999). As a result, the synaptic properties of neurons in the central nervous system have remained largely unexplored.

Several reporters of synaptic activity that are suitable for *in vivo* analysis are based on pHluorin, a highly pH-sensitive variant of the green fluorescent protein (Miesenböck et al., 1998). pHluorin and its derivatives are minimally fluorescent at the acidic pH conditions characteristic of the synaptic vesicle lumen, but highly fluorescence at neutral extracellular pH. As a result, synaptic vesicle exocytosis results in a sharp increase in the fluorescence of pHluorin fusion proteins targeted to the synaptic vesicle lumen. Their subsequent endocytosis and re-acidification quenches fluorescence, providing readouts at multiple stages of the synaptic vesicle cycle (Di Giovanni and Sheng, 2015; Fernandez-Alfonso and Ryan, 2008; Li et al., 2005; Sankaranarayanan and Ryan, 2000).

The genetic tractability and transparency of *C. elegans* are ideal for pHluorin imaging, and indeed, pHlourin-synaptobrevin fusion proteins have been used to study steady-state synaptic properties at the neuromuscular junction and in several other neurons (Dittman and Kaplan, 2006; Oda et al., 2011; Voglis and Tavernarakis, 2008). However, the standing plasma membrane levels of pHlourin-synaptobrevin fusion proteins make them ill-suited to real-time analysis (Dittman and Kaplan, 2006). By contrast, the vesicular glutamate transporter (VGLUT) has a minimal residence time on the plasma membrane in mammalian neurons (Voglmaier et al., 2006), and therefore is better suited for pHluorin imaging of vesicle dynamics (Balaji and Ryan, 2007).

We show here that EAT-4 VGLUT-pHlourin fusions can be used to study dynamic release and retrieval of synaptic vesicles from individual neurons in intact *C. elegans*. Using VGLUT-pHluorin fusions, we show that the release and retrieval of glutamatergic synaptic vesicles in two sensory neurons, AWC^ON^ and ASH, are kinetically distinct and matched to their signaling properties. We further demonstrate differential contributions of SNARE regulators to synaptic dynamics in AWC^ON^ and ASH, and describe activity-dependent regulation of AWC^ON^ exo- and endocytosis.

## Results

### VGLUT-pHluorin reports synaptic activity in AWC^ON^ and ASH neurons

The AWC^ON^ and ASH sensory neurons, which sense attractive odors and repulsive chemical and physical stimuli, respectively, are dynamically and molecularly distinct (Bargmann, 2006). Based on calcium imaging studies, the AWC^ON^ olfactory neurons are tonically active at rest, inhibited (likely hyperpolarized) by odor stimuli, and transiently activated upon odor removal before a return to baseline (Chalasani et al., 2007)(Figure 1A). This pattern resembles that of vertebrate photoreceptors, and indeed these neurons utilize a similar cGMP sensory transduction cascade (Bargmann, 2006; Zhang and Cote, 2005). By contrast, calcium imaging and electrophysiological studies of the ASH nociceptive neurons indicate that they are strongly activated by noxious chemical and mechanical stimuli, and recover quickly upon stimulus removal (Hilliard et al., 2005; Geffeney et al., 2011)(Figure 1B). Like vertebrate nociceptive neurons, ASH neurons signal via G protein-regulated TRPV channels (Bargmann, 2006). Both AWC^ON^ and ASH signal to downstream neurons through glutamatergic synapses and the vesicular glutamate transporter EAT-4 (Chalasani et al., 2007; Lee et al., 1999).

**Figure 1.**
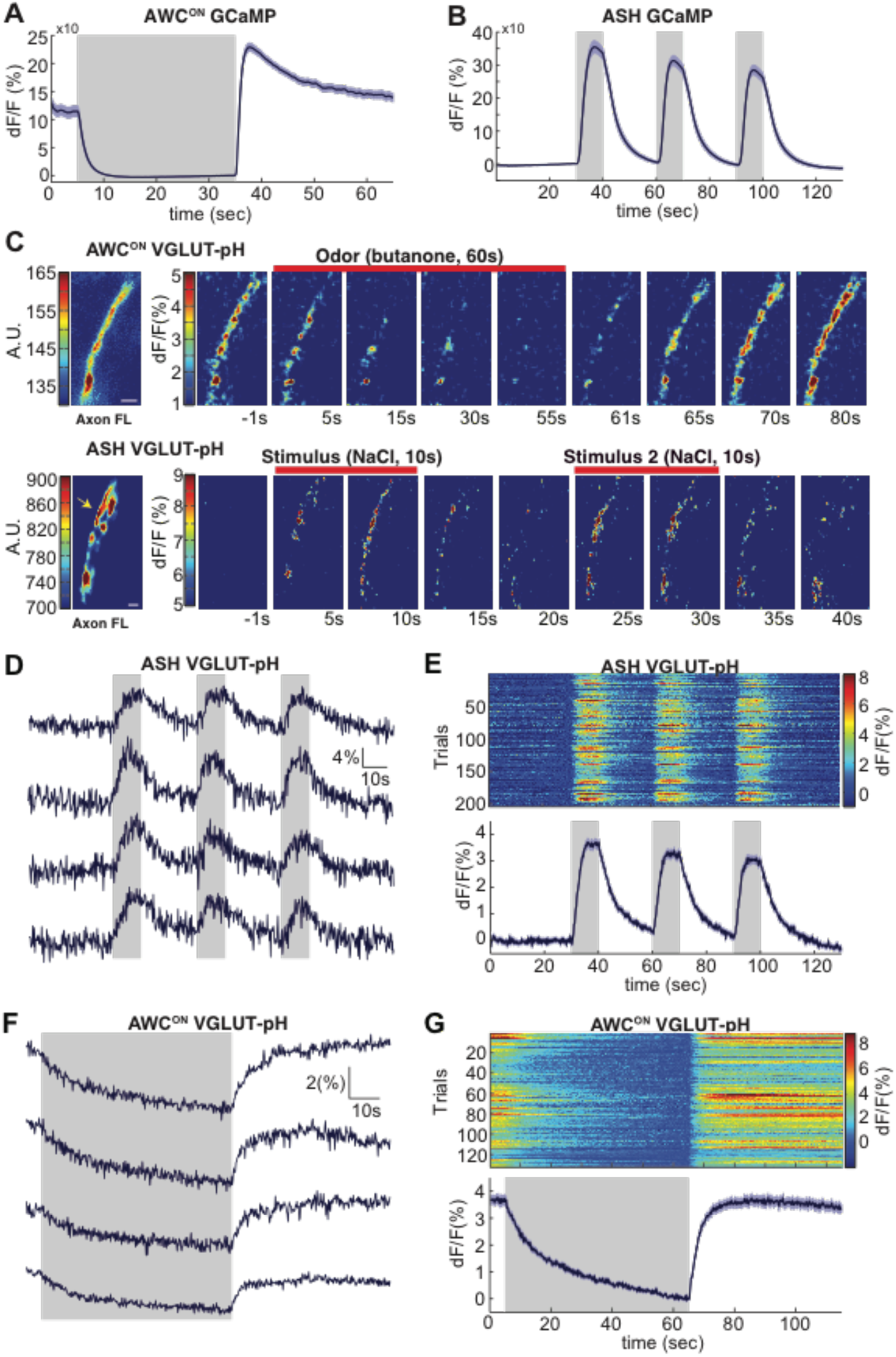
Sensory stimuli evoke VGLUT-pH signals in AWC^ON^ and ASH neurons. **(A)** AWC^ON^ GCaMP5A responses measured at the cell body in response to butanone stimulation (n=47, 16 animals, 2-3 trials each). **(B)** ASH GCaMP3 responses measured at the cell body in response to 500 mM NaCl stimulation (n=39, 13 animals, 3 trials each). **(C)** Individual AWC^ON^ and ASH VGLUT-pH responses. Left: Fluorescence intensity of VGLUT-pH along the axon prior to stimulation (a.u. arbitrary units, converted into reference F value in right panels). White scale bars = 2 um. Right: Images of VGLUT-pH fluorescence changes upon butanone removal (AWC^ON^) or NaCl addition (ASH) at t=0. Recordings were smoothed using a running average in time and space (3 frames, 3 pixels in x and y). **(D,F)** Single trials of ASH and AWC^ON^ VGLUT-pH responses from four individuals. Top trace in each panel is from the axon in (C). **(E,G)** Population ASH and AWC^ON^ VGLUT-pH responses. Top panel: Heat map of individual trials, 3 per animal, presented in sequential order. Bottom panel: Mean response from all trials. AWC^ON^: 132 trials from 44 animals, 3 trials each. ASH: 204 trials from 68 animals, 3 trials each. Gray bars mark stimulus periods. Blue shading indicates S.E.M.

To image synaptic vesicle (SV) endo- and exocytosis from single neurons in intact animals, we inserted super-ecliptic pHluorin into the first lumenal domain of EAT-4 (VGLUT-pH) and expressed this fusion protein using cell-specific promoters for AWC^ON^ and ASH (Fig S1A-C). Immobilized animals were imaged in microfluidic chips that enable the precise delivery and removal of chemical stimuli and simultaneous monitoring of cell fluorescence at high magnification (Chalasani et al., 2007; Chronis et al., 2007). AWC^ON^ responses were elicited by addition and removal of butanone odor, while ASH responses were elicited by addition and removal of a noxious 0.5 M NaCl stimulus, under conditions that gave robust signals with GCaMP calcium sensors (Fig 1A,B).

The VGLUT-pH reporter was exclusively localized to the axon in both AWC^ON^ and ASH, with a semi-punctate distribution in synaptic regions (Fig 1C, left panels). Delivery of butanone to AWC^ON^ resulted in a reduction of VGLUT-pH fluorescence with recovery after odor removal (Fig 1C, top). Delivery of NaCl to ASH resulted in an increase in VGLUT-pH fluorescence, followed by a decrease after NaCl removal (Fig 1C, bottom). In both neurons, responses were observed across the ∼20 um region of the axon that was imaged, and could be followed with single trial resolution (Fig 1D-G & Supplemental Methods). AWC^ON^ and ASH synapses appeared to be highly reliable, as virtually every stimulus triggered a detectable VGLUT-pH response (Fig 1E,G).

In ASH, VGLUT-pH fluorescence rose rapidly upon stimulus addition, and fell immediately upon stimulus removal, closely resembling the calcium response (Fig 1D,E, compare 1B). These results are consistent with a model in which ASH synaptic vesicle exocytosis is induced by stimulus-triggered calcium entry, and terminates rapidly with subsequent endocytosis and reacidification. In a control experiment, pHluorin tethered to the extracellular face of the ASH plasma membrane as a CD4 fusion protein did not respond to NaCl with fluorescence changes (Fig S1D-E), indicating that the signal reflects synaptic vesicle dynamics and not changes in extracellular pH.

In AWC^ON^, VGLUT-pH fluorescence decreased throughout a one minute odor presentation, and odor removal resulted in a rapid increase to pre-stimulus levels without an overshoot (Fig 1F,G). The properties are consistent with a kinetic model in which the AWC^ON^ neuron has tonic synaptic vesicle release, with basal VGLUT-pH fluorescence determined by steady-state levels of exocytosis versus endocytosis and reacidification. In this model, odor addition reduces calcium and synaptic vesicle exocytosis, and odor removal triggers a calcium increase, synaptic vesicle exocytosis, and a return to the pre-stimulus steady state.

A close examination of VGLUT-pH signals after odor removal showed that AWC^ON^ fluorescence had an initial fast rate of increase for ∼1s, and then transitioned to a slower rate of increase over the following ∼5s (Fig 2A,B, Fig S2A). These dynamics suggest that synaptic vesicle exocytosis is transiently enhanced above its basal level immediately after odor removal, in agreement with the known calcium overshoot in the AWC^ON^ cell body (e.g. Fig 1A). To better understand the relationship between somatic calcium, synaptic calcium, and exocytosis in AWC^ON^, we fused a GCaMP calcium sensor to the synaptic vesicle protein synaptogyrin (Fig S1C). This protein labeled the axons in a punctate pattern consistent with synapses, and should detect calcium levels in the immediate vicinity of synaptic vesicles. Synaptic calcium levels monitored with synaptogyrin-GCaMP rose and fell substantially more quickly than those in the cell body (Fig 2C,D; compare Fig 1A). The peak rate of calcium entry, at ∼1s after odor removal, occurred at the same time as the peak rate of VGLUT-pH fluorescence increase (Fig 2B,D,G).

**Figure 2.**
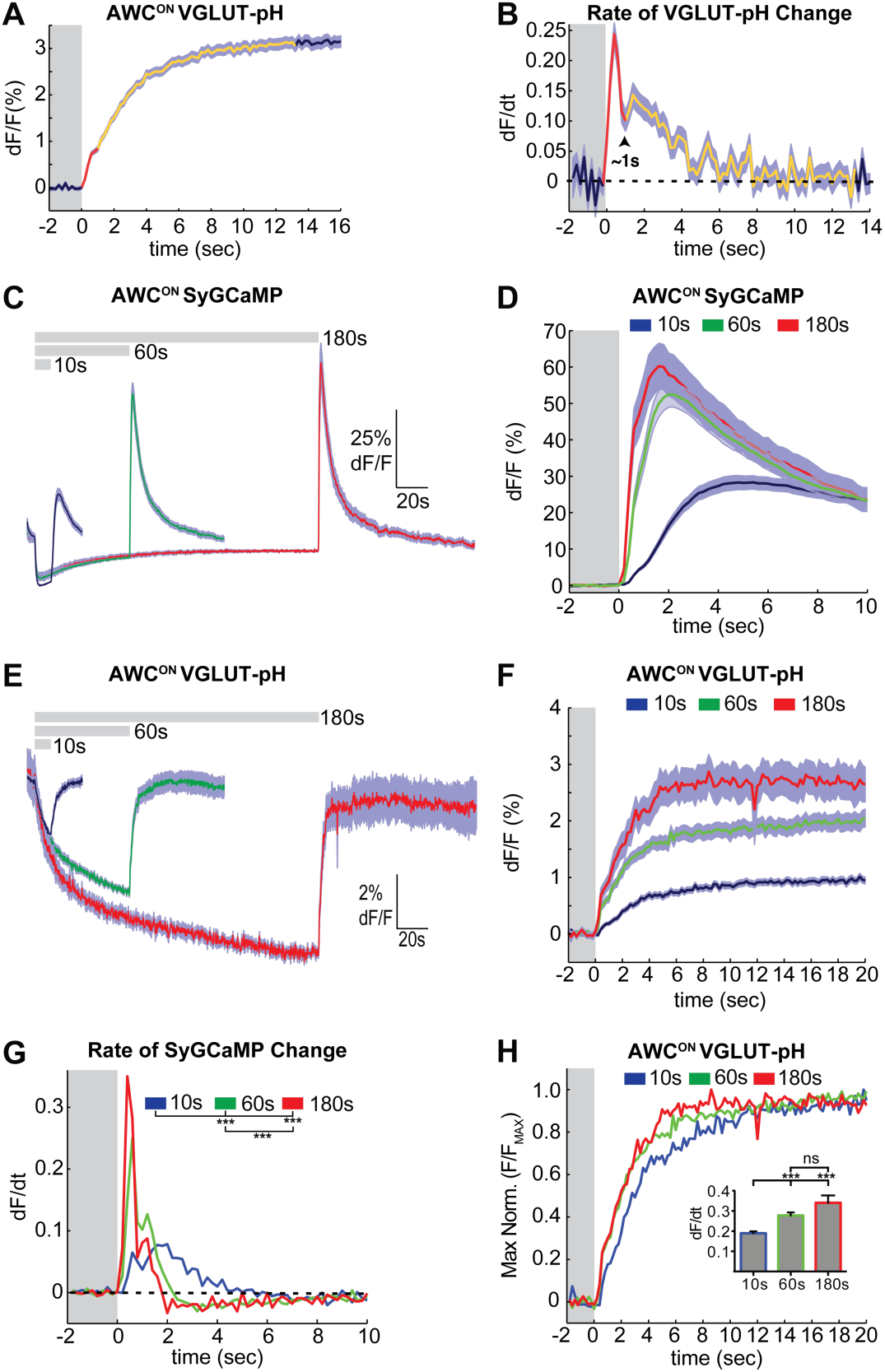
Two kinetic phases of VGLUT-pH responses and calcium influx in AWC^ON^. **(A)** Mean AWC^ON^ Vglut-pH signal upon odor removal (60s stimulus, n=219 trials, 1-3 trials per animal). Trace is colored according to transitions in time derivative in (B). **(B)** Mean time derivative of AWC^ON^ VGLUT-pH signals in (A). **(C)** AWC^ON^ synaptic calcium responses to odor measured with syGCaMP; traces aligned to odor addition. 10s pulses: n = 42 (7 animals, 6 trials each). 60s pulses: n = 21 (7 animals, 3 trials each). 3 min pulses: n = 7 (7 animals, 1 trial each). **(D)** syGCaMP responses from (C) aligned to odor removal. **(E)** AWC^ON^ VGLUT-pH responses to butanone pulses; traces aligned to odor addition. 10s pulses: n = 120 (20 animals, 6 trials each). 60s pulses: n = 59 (21 animals, 2-3 trials each). 3 min pulses: n = 14 (14 animals, 1 trial each). **(F)** VGLUT-pH responses from (E) aligned to odor removal. **(G)** Mean time derivative of AWC^ON^ syGCaMP signals in (D) shows different peak rates 1s after odor removal (*** p < 0.0001, one-way ANOVA with Tukey’s correction). **(H)** Average VGLUT-pH odor removal responses from (F) after normalizing response magnitude. Inset: Average peak time derivative of AWC^ON^ VGLUT-pH signals 1s after odor removal. *** p < 0.0001, ns (p =0.14), One-way ANOVA with Tukey’s correction. For time derivative plots each individual trial was smoothed with a running average (3 frames) before taking the derivative. Units are change in dF/F (%) per 200ms. Gray bars in A,B,D,F-H mark odor stimulus periods. Shading indicates S.E.M.

Varying the duration of odor exposure prior to odor removal allowed a more focused comparison of synaptic calcium dynamics and VGLUT-pH dynamics in AWC^ON^ (Fig 2C-F). Removing odor after a 10s exposure elicited a small synaptic calcium overshoot within the first second (Fig 2D,G), and a small increase in exocytosis during the same interval (Fig 2H). Both synaptic calcium and exocytosis rates were elevated more substantially for ∼1s after a 60s or 180s odor exposure (Fig 2F-H). This correspondence suggests that the transient synaptic calcium overshoot following long odor stimuli evokes a brief pulse of synaptic vesicle release above the tonic level.

### The SNARE complex drives tonic and evoked synaptic vesicle release

Synaptic vesicle release is triggered by the SNARE complex, which is composed of the plasma membrane proteins syntaxin and SNAP-25, and the vesicle-associated protein synaptobrevin. Expressing the light chain of tetanus toxin (TeTx), which cleaves synaptobrevin (Schiavo et al., 1992), in either AWC^ON^ or ASH eliminated their VGLUT-pH responses, as predicted if the VGLUT-pH signals report synaptic vesicle dynamics (Fig 3A,B). Notably, AWC^ON^ VGLUT-pH fluorescence did not decrease upon odor addition in TeTx animals, suggesting that tonic AWC^ON^ exocytosis as well as the evoked exocytosis after long odor stimuli requires the SNARE complex.

**Figure 3.**
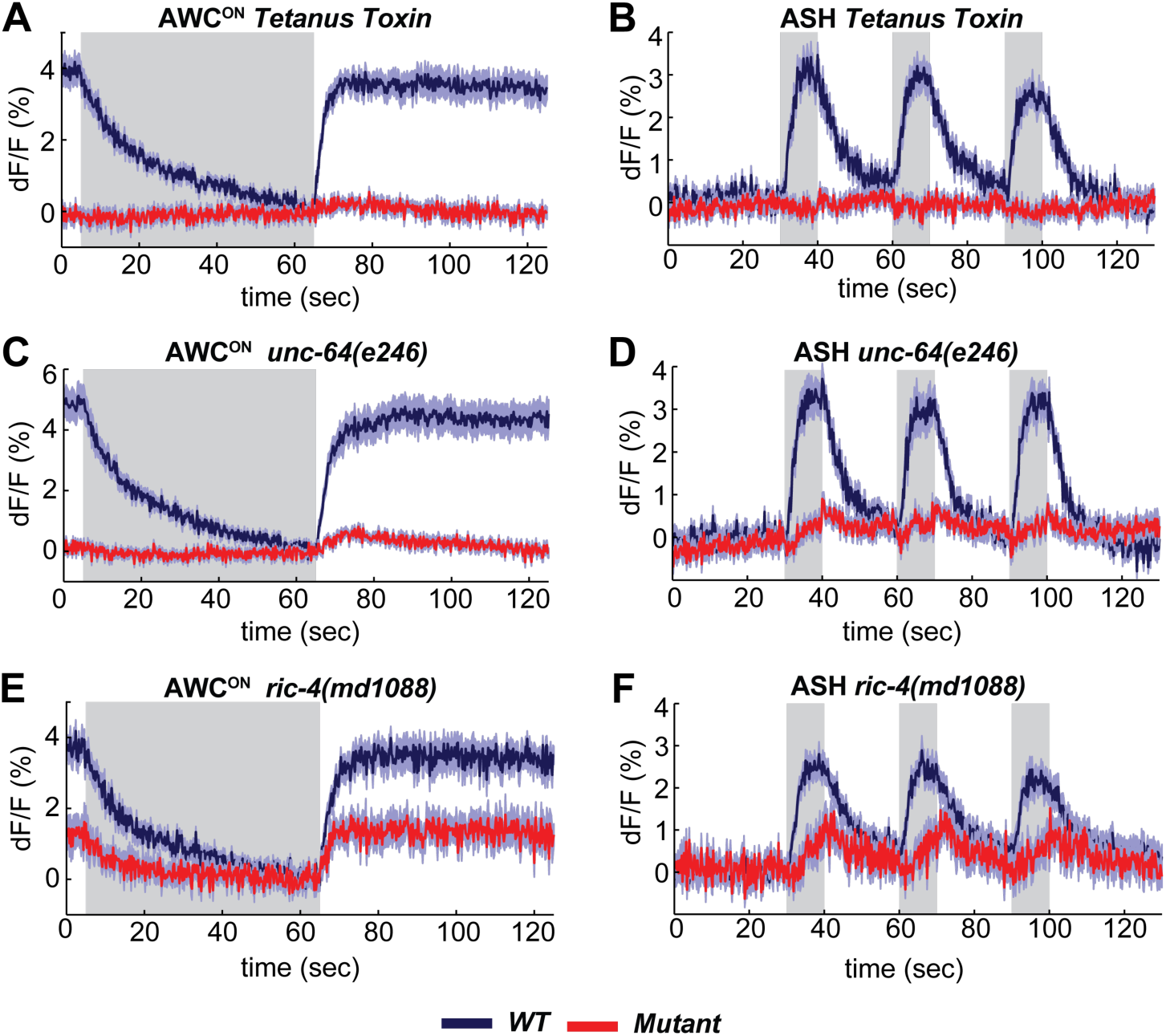
The SNARE complex is required for VGLUT-pH responses in AWC^ON^ and ASH. Wild-type controls (blue) and mutants (red) measured in parallel. **(A,B)** VGLUT-pH responses are eliminated by cell-specific expression of TeTx light chain. (A) AWC^ON^ *str-2* promoter driving TeTx n=25 (9 animals, 2-3 trials each). AWC^ON^ *wt* (TeTx-array negative animals tested in parallel) n=28 (10 animals, 2-3 trials each). (B) ASH *sra-6* promoter driving TeTx n=33 (11 animals, 3 trials each). ASH *wt* (TeTx-array negative animals tested in parallel) n=24 (8 animals, 3 trials each). **(C,D)** VGLUT-pH responses in *unc-64(e246)* (partial loss of function) syntaxin mutants. (C) AWC^ON^ *unc-64(e246)* n=30 (10 animals, 3 trials each). AWC^ON^ *wt* n=23 (9 animals, 2-3 trials each). (D) ASH *unc-64(e246)* n=24 (8 animals, 3 trials each). ASH *wt* n=18 (6 animals, 3 trials each). **(E, F)** VGLUT-pH responses in *ric-4(md1088)* (partial loss of function) SNAP-25 mutants. (E) AWC^ON^ *ric-4(md1088)* n=12 (4 animals, 3 trials each). AWC^ON^ *wt* n=15 (5 animals, 3 trials each). (F) ASH *ric-4(md1088)* n=16 (6 animals, 2-3 trials each). ASH *wt* n=32 (11 animals, 2-3 trials each). The molecular nature of mutations is described in Supplemental Methods.,All differences are significant (p<0.0001, unpaired t-test), as detailed in Supplemental Table 1. Gray bars mark stimulus periods. Shading indicates S.E.M.

Mutant analysis supported the roles of SNARE complex proteins in sensory exocytosis. Null mutations in SNARE complex proteins are inviable, but partial loss of function in syntaxin (*unc-64*) and SNAP-25 (*ric-4)* are viable, with reduced synaptic vesicle release at the *C. elegans* neuromuscular junction (Ogawa et al., 1998; Staunton et al., 2001). Both *unc-64(e246)* (Fig 3C,D) and *ric-4(md1088)* (Fig 3E,F) had diminished VGLUT-pH responses in AWC^ON^ and ASH. These results are consistent with a requirement for the SNARE complex in tonic and evoked glutamate release from AWC^ON^ and evoked glutamate release from ASH.

### SNARE regulators can differentially affect AWC^ON^ and ASH

A suite of conserved presynaptic proteins including SNARE-associated proteins, scaffold proteins, and small GTPases affect synaptic release in many animals, but their apparent importance varies between reports. Among these presynaptic regulators are UNC-13 and UNC-18 (Augustin et al., 1999; Varoqueaux et al., 2002; Verhage et al., 2000). UNC-13 is implicated in priming synaptic vesicles prior to release; it is a multidomain protein with a MUN domain that can open a closed conformation of syntaxin, and three C2 domains, which bind phorbol esters, phospholipids, and in some cases calcium (Richmond et al., 1999; Michelassi et al., 2017). ASH neurons had no detectable VGLUT-pH response to sensory stimuli in *unc-13* null mutants, indicating an absolute requirement for this protein in ASH synaptic vesicle mobilization (Fig 4B). By contrast, AWC^ON^ neurons in *unc-13* null mutants had a significant, albeit reduced, increase in VGLUT-pH signal after odor removal indicative of residual synaptic activity (Fig 4A).

**Figure 4.**
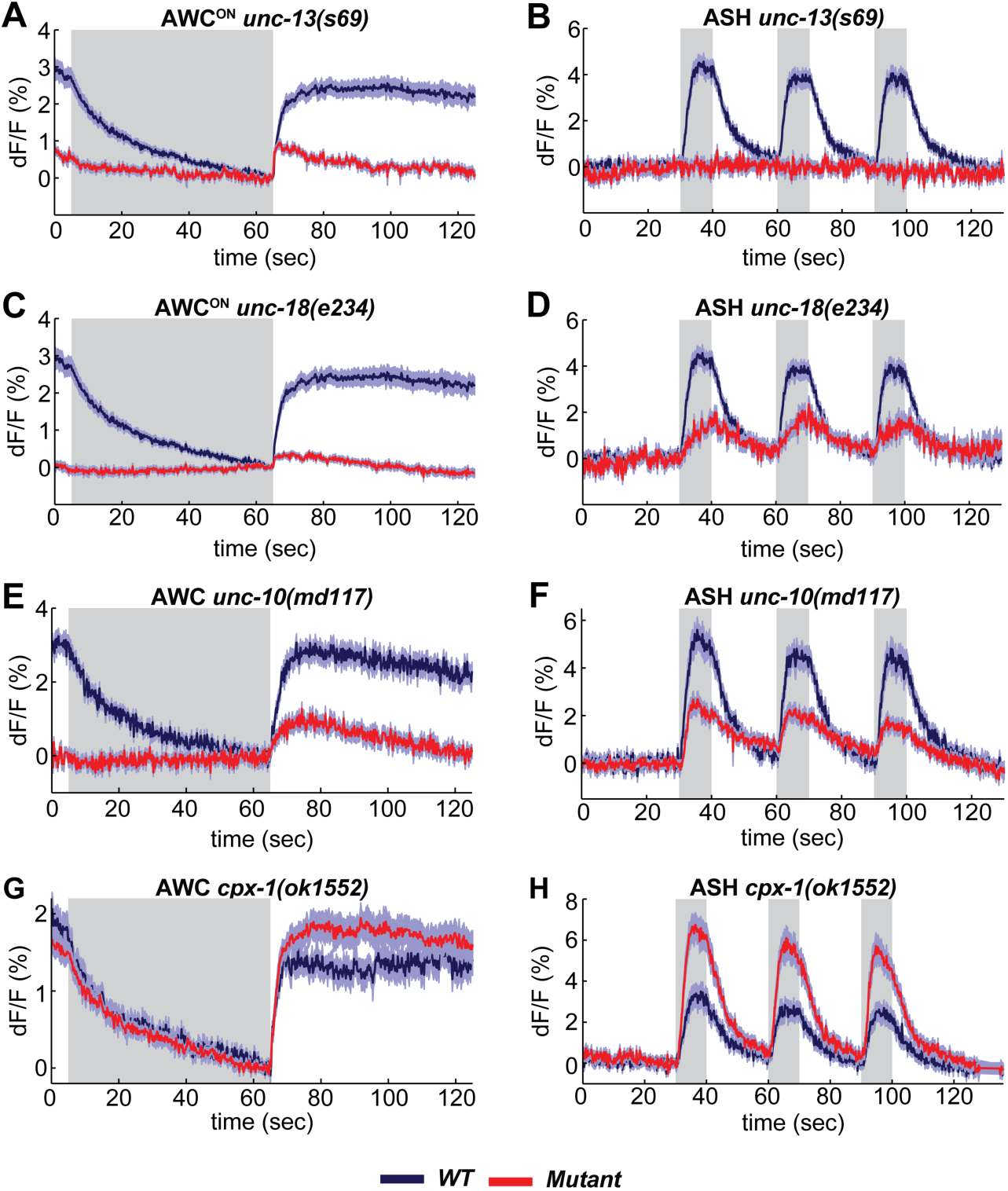
SNARE regulators differentially affect AWC^ON^ and ASH. Wild-type controls (blue) and mutants (red) measured in parallel. **(A,B)** VGLUT-pH responses in *unc-13(s69)* null mutants. (A) AWC^ON^ responses in *unc-13(s69),* n=25 (9 animals, 2-3 trials each) and *wt,* n=33 (11 animals, 3 trials each). (B) ASH responses in *unc-13(s69),* n=18 (7 animals, 1-3 trials each) and *wt,* n=42 (14 animals, 3 trials each). **(C,D)** VGLUT-pH responses in *unc-18(e234)* mutants. (C) AWC^ON^ responses in *unc-18(e234),* n=25 (9 animals, 1-3 trials each) and *wt,* n=33 (11 animals, 3 trials each). (D) ASH responses in *unc-18(e234),* n=22 (8 animals, 1-3 trial each) and *wt,* n=42 (14 animals, 3 trials each). **(E,F)** VGLUT-pH responses in *unc-10(md117)* mutants. (E) AWC^ON^ responses in *unc-10(md117),* n=23 (8 animals, 1-3 trials each), and *wt,* n=27 (9 animals, 3 trials each). (F) ASH responses in *unc-10(md117),* n=33 (12 animals, 2-3 trials each), and *wt,* n=24 (8 animals, 3 trials each). **(G,H)** VGLUT-pH responses in *cpx-1(ok1552)* mutants. (G) AWC^ON^ responses in *cpx-1(ok1552),* n=36 (12 animals, 3 trials each), and *wt,* n=17 (6 animals, 2-3 trials each). (H) ASH responses in *cpx-1(ok1552),* n=38 (13 animals, 2-3 trials each), and *wt* n=33 (11 animals, 3 trials each). WT and mutants are significantly different in panels A-F and H, as detailed in Supplemental Table 2. Gray bars mark stimulus periods. Shading indicates S.E.M.

UNC-18 also interacts with syntaxin, and regulates syntaxin localization as well as activity (McEwen and Kaplan, 2008; Ogawa et al., 1998). AWC^ON^ VGLUT-pH responses were nearly eliminated in *unc-18* null mutants (Fig 4C). However, *unc-18* ASH neurons responded to stimuli by mobilizing VGLUT-pH, albeit to a lesser extent than the wild-type (Fig 4D). These results reveal heterogeneity in the requirements for SNARE regulators in different cell types: ASH has a stronger requirement for *unc-13* and a weaker requirement for *unc-18* than AWC^ON^.

Other SNARE regulators had similar effects on ASH and AWC^ON^. In both neurons, synaptic vesicle release was reduced but not eliminated by *unc-10/RIM* mutations (Fig 4E,F). Similarly, both ASH and AWC^ON^ showed enhanced synaptic vesicle release in *cpx-1/complexin* mutants (Fig 4G-H), although this effect was only statistically significant in ASH (Table S2).

### Cytoplasmic pH is increased by neuronal activity, independent of synaptic release

As a counterpoint to the VGLUT-pHluorin experiments, we examined the effect of sensory stimuli on cytoplasmic pH in AWC^ON^ and ASH neurons. Activity-dependent increases or decreases in cytoplasmic pH have been documented in both vertebrate and invertebrate neurons (Chesler, 2003; Rossano et al., 2013; Zhang et al., 2010). Similarly, expressing an untagged super-ecliptic pHluorin in the cytoplasm of AWC^ON^ and ASH neurons (cyto-pH) reported robust stimulus-dependent pH changes. Odor addition increased cyto-pH fluorescence in AWC^ON^, suggesting alkalinization, and odor removal decreased cyto-pH fluorescence, suggesting activity-dependent acidification (Fig 5A). These pH changes were opposite in sign to the signals defected by VGLUT-pH (compare Fig 5A to 1G), and were strongest in the axon, intermediate in the cell body, and weak in the sensory dendrite (Fig 5A). In ASH, NaCl stimuli elicited a decrease in cyto-pH fluorescence suggestive of acidification, again the opposite sign of the VGLUT-pH signal (Fig 5B).

**Figure 5.**
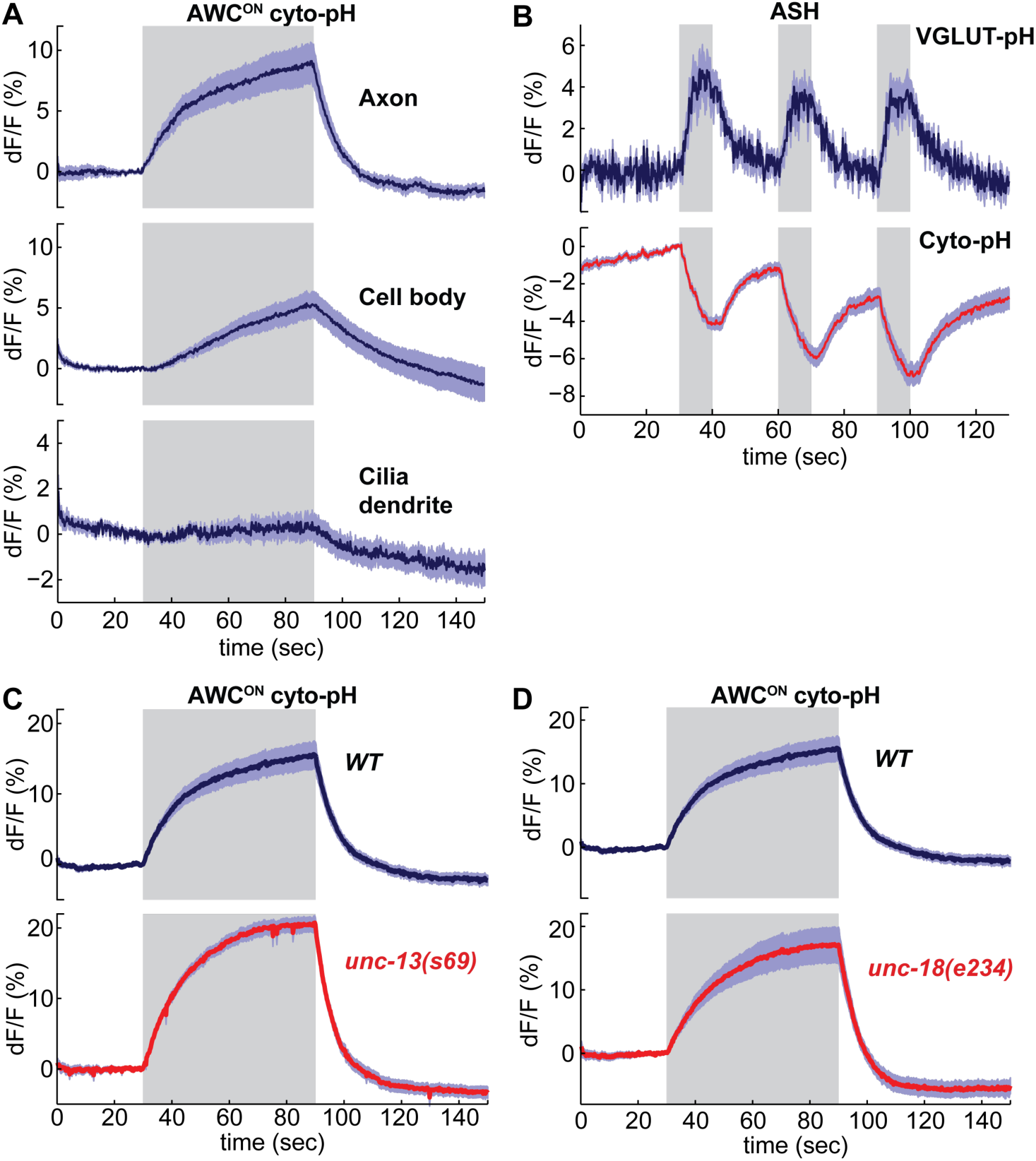
Sensory stimuli evoke cytoplasmic pH changes in AWC^ON^ and ASH. **(A)** Average cyto-pH signals at different subcellular sites of AWC^ON^. Axon and cell body, n=9 (3 animals, 3 trials each), Cilia-dendrite n=6 (2 animals, three trials each). **(B)** Average ASH VGLUT-pH response (top) and cyto-pH response (bottom), tested in parallel under the same stimulus conditions. VGLUT-pH n=9 (3 animals, 3 trials each). cyto-pH n=27 (9 animals, 3 trials each). **(C,D)** Average AWC^ON^ cyto-pH responses in (C) *unc-13(s69)* and (D) *unc-18(e234)* mutants. Mutants and wild-type controls were measured on the same days; neither mutant was significantly different from wild-type (p > 0.12, one-way ANOVA with Tukey’s correction). *unc-13(s69)* n=15 (5 animals, 3 trials each). *unc-18(e234)* n=11 (4 animals, 2-3 trials each). *wt* n=18 (6 animals, 3 trials each). Gray bars mark stimulus periods. Shading indicates S.E.M.

Unlike VGLUT-pH responses, AWC^ON^ cyto-pH signals were normal in *unc-13* or *unc-18* mutations (Fig 5C,D). These results suggest that cytoplasmic pH changes are independent of synaptic vesicle release.

### The endocytosis-reacidification process is accelerated by AP180/pCALM

The decrease in VGLUT-pH fluorescence at synapses represents the recapture of the protein from the cell surface by endocytosis and the acidification of the resulting synaptic vesicles (Balaji and Ryan, 2007; Sankaranarayanan and Ryan, 2000)(Fig S1C). To estimate the rate of this combined retrieval step, we fit fluorescence decreases from individual AWC^ON^ and ASH trials to a single exponential decay function (Fig 6A,B) (Smith et al., 2008). A large fraction of traces were consistent with single exponential decay, with an 18s time constant for AWC^ON^ and an 8s time constant for ASH (Fig 6C,D); a subset of traces were consistent with double exponential decay (Fig S3). The measured decay constants within and between neurons did not correlate with axon fluorescence before endocytosis (Fig S4), suggesting that VGLUT-pH expression levels did not saturate the endocytosis machinery (Sankaranarayanan and Ryan, 2000).

**Figure 6.**
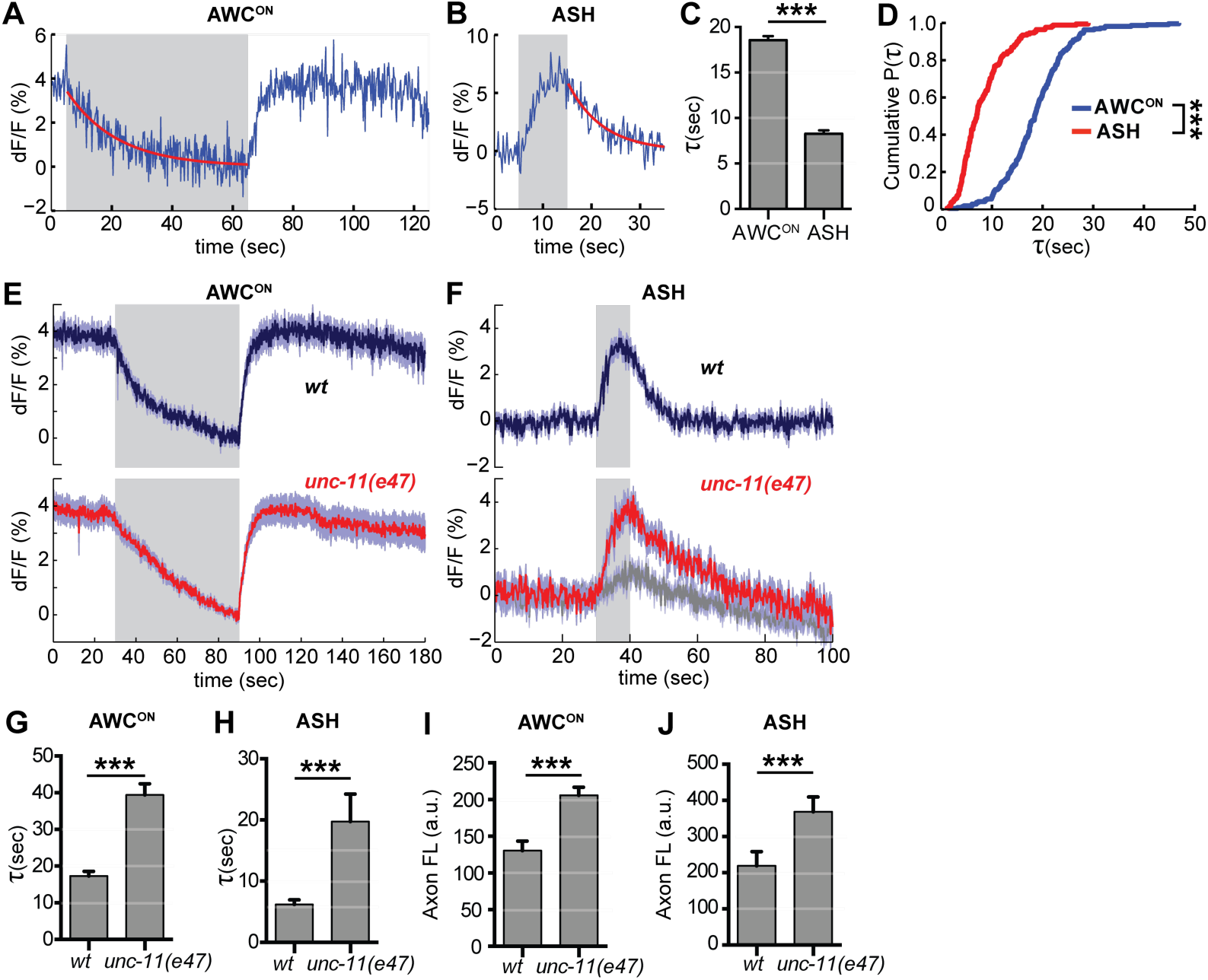
Synaptic vesicle retrieval is accelerated by AP180/CALM. **(A,B)** Representative single exponential fits (red) to single trials of (A) AWC^ON^ and (B) ASH VGLUT-pH decays upon stimulus addition or removal, respectively. For each neuron, some responses were consistent with double exponential decay models (Fig S3). **(C)** Average time constant of AWC^ON^ and ASH decays from single exponential fits. AWC^ON^ n=218 (76 animals, 2-3 trials each). ASH n=168 (56 animals, 2-3 trials each). *** p < 0.0001, unpaired t-test. **(D)** Empirical cumulative distribution plot of data in (C). Distributions differ by Kolmogorov-Smirnov test, *** p<0.0001. **(E,F)** Average VGLUT-pH responses in *unc-11(e47)* mutants. (E) AWC^ON^ responses in *unc-11(e47)* n=27 (10 animals, 2-3 trials each), *wt* n=15 (5 animals, 3 trials each). One *unc-11(e47)* animal did not respond and was removed from the analysis. (F) ASH responses in *unc-11(e47)* mutants. *wt* n=21 (7 animals, 3 trials each). *unc-11(e47)* n=23 (8 animals, 2-3 trials each). Red trace: mean of 9 trials (5 animals, 1-2 trails each) with clear responses to odor addition. Magnitude of response does not differ from *wt* (p=0.72, peak odor response); Gray trace: mean of 14 trials that produced weak or non-detectable responses to odor addition, significantly different from *wt* (p < 0.0001, peak odor response). One-way ANOVA, Tukey’s correction. **(G,H)** Average time constants from single exponential fits (initial 20s of decay) of data in (E,F). For ASH *unc-11(e47)* mutants, only data from the red trace was used. Unpaired t-test, p < 0.0001. AWC^ON^ *unc-11(e47)* n=25 (10 animals, 2-3 trials each); *wt* n=15 (5 animals, 3 trials each). ASH n as in (F). **(I,J)** Average axon fluorescence (first 5 frames of the recording). (I) AWC^ON^ *wt* n=12 animals, *unc-11(e47)* n=19 animals. (J) ASH *wt* n=7 animals, *unc-11(e47)* n=8 animals. Unpaired t-test, p < 0.0001. Gray bars mark stimulus periods. Shading and error bars indicate S.E.M.

Among the proteins most strongly implicated in synaptic vesicle retrieval is the adaptor protein AP180, which clusters synaptic vesicle proteins (Gimber et al., 2015; Koo et al., 2011) and interacts with AP-2/clathrin at a sorting stage immediately after endocytosis (Koo et al., 2015). The *C. elegans* AP180/CALM homolog *unc-11* has long been proposed to affect endocytosis, as well as affecting synaptic vesicle morphology and protein sorting (Nonet et al., 1999). Indeed, in AWC^ON^ neurons upon odor addition, and in ASH neurons after stimulus removal, *unc-11(e47)* null mutants had significantly slowed VGLUT-pH retrieval (Fig 6E-H). Exocytosis may also be impacted in *unc-11(e47)* mutants, as a significant fraction of trials produced weak or undetectable responses in ASH (Fig 6F, gray trace). Baseline VGLUT-pH fluorescence was higher in *unc-11(e47)* mutants than in wild type for both neurons, consistent with increased VGLUT-pH on the cell surface or in other neutral compartments (Fig 6I,J).

### Activity-dependent regulation of VGLUT-pH retrieval in AWC^ON^

The distinct endocytosis and recapture rates in AWC^ON^ and ASH could reflect either cell type-specific or cell state-specific processes. With respect to cell state, GCaMP measurements in AWC^ON^ and ASH suggest that the calcium levels in ASH at the end of a NaCl stimulus are higher than basal calcium levels in AWC^ON^ when odor is added (Fig 1A,B). This difference in calcium levels could affect vesicle traffic, as endocytosis in other systems is accelerated at high calcium concentrations (Leitz and Kavalali, 2016; Neves et al., 2001; Sankaranarayanan and Ryan, 2001). To separate the effects of cell type and cell state, we compared VGLUT-pH retrieval in AWC^ON^ at basal and elevated calcium levels. Odor was delivered to AWC^ON^, removed after one minute to elicit a calcium overshoot, and then delivered again after 10s while calcium levels were still elevated (Fig 7A). Strikingly, AWC^ON^ VGLUT-pH retrieval was accelerated during the second odor pulse, matching the ∼8s retrieval time observed in ASH (Fig 7B,C). The effect was temporary (lasting < 70 seconds) but could be induced again after another 60s odor pulse (Fig 7D). *unc-11(e47)* mutants were also regulated by the dual odor-pulse protocol, and delayed compared to wild-type under both conditions (Fig 7E,F). These results indicate that VGLUT-pH retrieval in AWC^ON^ is regulated by activity, consistent with calcium-dependent acceleration of endocytosis.

**Figure 7.**
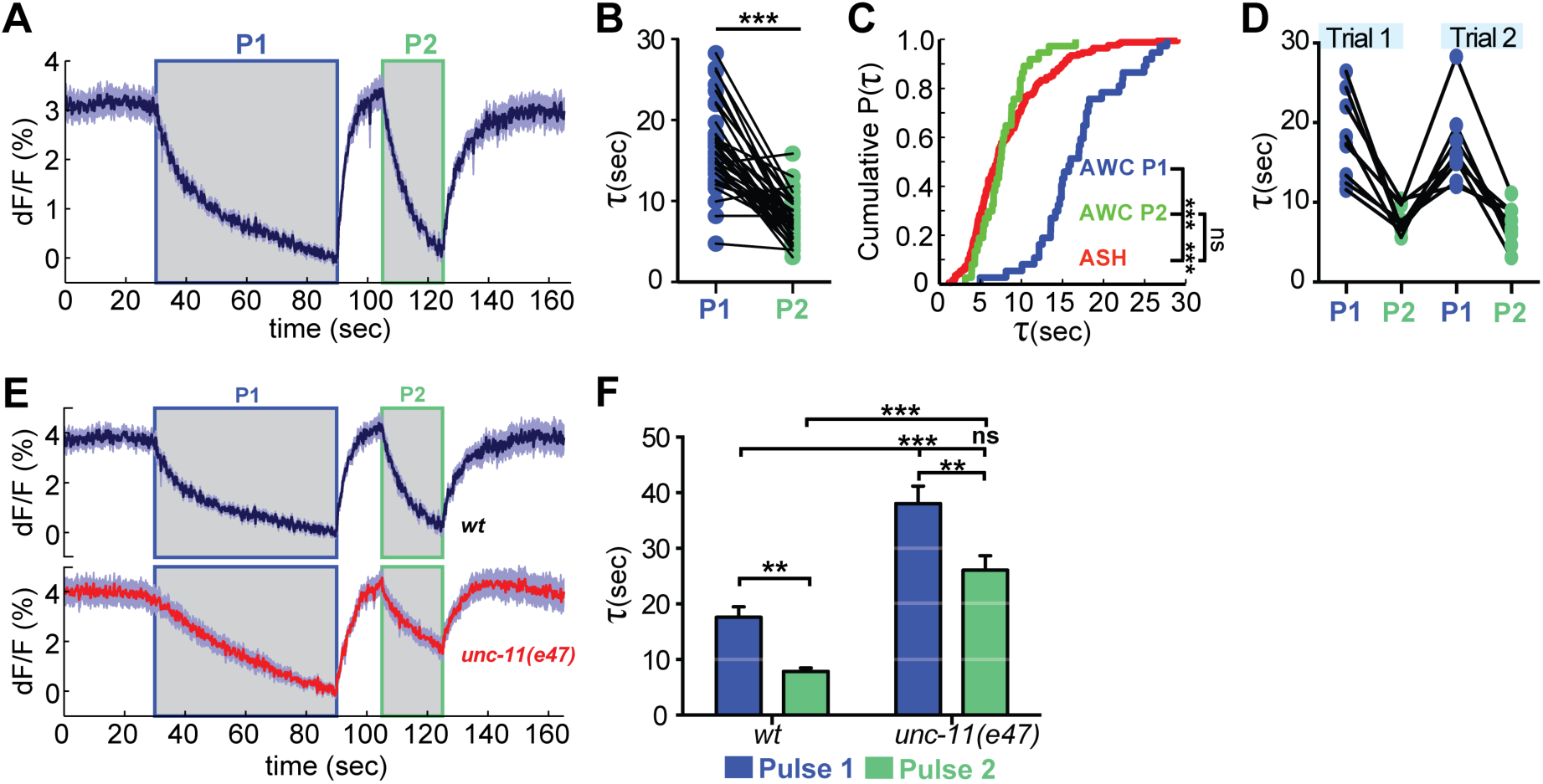
Recent neural activity modulates AWC^ON^ VGLUT-pH retrieval. **(A)** Average AWC^ON^ VGLUT-pH response to two successive odor stimuli, applied for 60s (P1) and 20s (P2). n=39 (13 animals, 3 trials each). **(B)** Time constants from single-term exponential fits to P1 and P2 from (A) performed on the initial 20s of the decay for each pulse. n=37 (13 animals, 2-3 trials each). Paired t-test, *** p<0.0001. **(C)** Cumulative distribution plot of time constants for AWC P1, AWC P2, and ASH VGLUT-pH decays. AWC P1 and AWC P2 data from (B) and ASH data from Fig 6D. Kruskal-Wallis & Dunn’s test for multiple comparisons, *** p< 0.0001, ns p = 0.1. **(D)** Time constants for P1 and P2 from two consecutive trials of the stimulation protocol in (A) (n=8 animals, 70s between trials). **(E)** Average AWC VGLUT-pH signals in *wt* and *unc-11(e47)* mutants. *wt* n=21 (7 animals, 3 trials each). *unc-11(e47)* n=25 (9 animals, 3 trials each) (two non-responding trials removed). **(F)** Average time constants from single exponential fits (initial 20s of decay) of data in (E). *wt* n=21 (7 animals, 3 trials each). *unc-11(e47)* n=22 (8 animals, 2-3 trials each). Two-way ANOVA, *** p<0.0001, ** p<0.008, ns (p = 0.07). Gray bars marks stimulus periods. Shading and error bars indicate S.E.M. ns = not significant.

### Protein kinase C epsilon regulates AWC^ON^ exocytosis downstream of calcium

In addition to demonstrating core neuronal properties, VGLUT-pH imaging provides a tool to better understand molecular regulators of synaptic transmission. For example, a protein kinase C epsilon (novel class) encoded by *pkc-1* is required for normal behavioral responses to odors detected by AWC^ON^, and has previously been suggested to act at a step downstream of AWC^ON^ calcium entry (Tsunozaki et al., 2008). VGLUT-pH imaging in AWC^ON^ can be used to examine this possibility in detail.

We confirmed the normal calcium response to odor in *pkc-1* mutants (Fig S5), and additionally determined that synaptic calcium signals detected with synaptogyrin-GCaMP were intact in *pkc-1* (Fig 8A,B, top). However, VGLUT-pH signals in *pkc-1* mutants were blunted and delayed (Fig 8A,B, bottom). These defects were comparable to those of strong SNARE mutations, and appeared to affect both the tonic release signal and the evoked release after removing short, intermediate, or long odor stimuli (Fig 8C).

**Figure 8.**
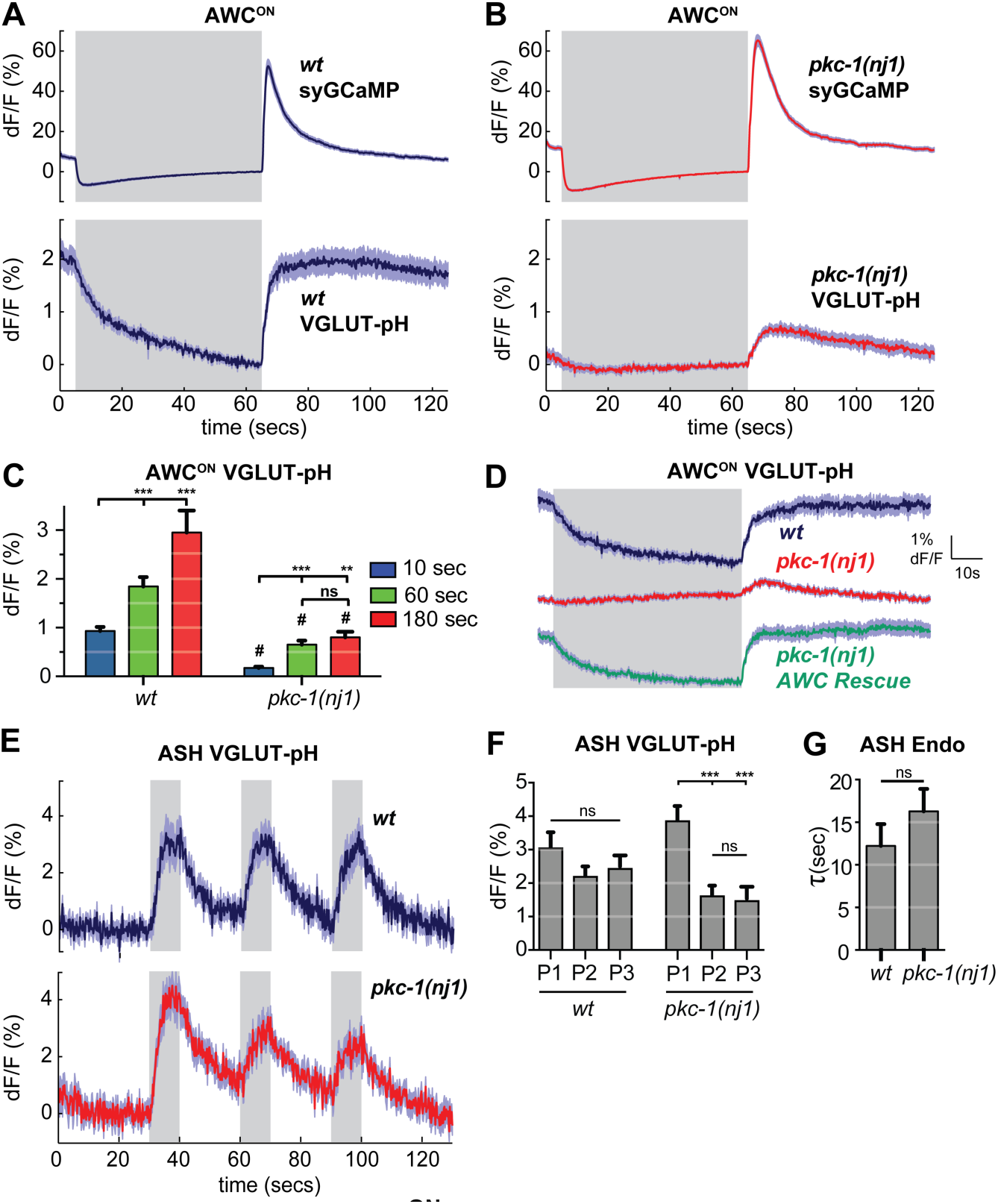
*pkc-1* regulates AWC^ON^ glutamate release downstream of calcium influx. **(A,B)** AWC^ON^ synaptic calcium (top) and VGLUT-pH (bottom) responses in (A) wild-type and (B) *pkc-1(nj1)* mutant animals. *wt* SyGCaMP n=21 (7 animals, 3 trials each), *pkc-1* SyGCaMP n=25 (9 animals, 1-3 trials each). *wt* VGLUT-pH n=29 (10 animals, 2-3 trials each), *pkc-1(nj1)* VGLUT-pH n=48 (17 animals, 2-3 trials each). **(C)** Average AWC VGLUT-pH peak response magnitude after odor removal for indicated stimulation durations. #, different from wild-type p <0.0001. ** p = 0.0026, *** p <0.0001. Two-way ANOVA Tukey’s correction. 10s pulses: *wt* n=60 (10 animals, 6 trials each). *pkc-1(nj1)* n=102 (17 animals, 4-6 trials each). 60s pulses: as in (A,B). 3 min pulses: *wt* n=8 (8 animals, 1 trial each). *pkc-1(nj1)* n=15 (15 animals, 1 trials each). **(D)** Expression of *pkc-1* cDNA in AWC^ON^ rescues *pkc-1(nj1)* VGLUT-pH responses. *wt* n=18 (6 animals, 3 trials each). *pkc-1(nj1)* n=30 (10 animals, 3 trials each). *pkc-1(nj1)* AWC^ON^ rescue n=30 (10 animals, 3 trials each). **(E)** ASH VGLUT-pH responses in *pkc-1(nj1)* mutants. *wt* n=15 (5 animals, 3 trials each). *pkc-1(nj1)* n=15 (6 animals, 2-3 trials each). **(F)** Average peak responses for each stimulus pulse (P1-3) within a trial. Data from (E). Two-way ANOVA Tukey’s correction. *** p<0.0001. **(G)** Average time constants from single exponential fits of data in (E, first pulse). Supporting statistical analysis is detailed in Supplemental Table 3 and Source Data. Gray bars mark stimulus periods. Shading and error bars indicate S.E.M. ns = not significant.

Selective transgenic expression of a *pkc-1* cDNA in AWC^ON^ resulted in full rescue of the VGLUT-pH defect (Fig 8D, Fig S5), indicating that the *pkc-1* defects in synaptic vesicle exocytosis were cell autonomous to AWC^ON^.

In contrast with AWC^ON^, VGLUT-pH signals in ASH neurons were only slightly affected by *pkc-1* mutations (Fig 8E-G); *pkc-1* mutants substantially preserved both exocytosis and retrieval dynamics in ASH. A subtle *pkc-1* defect was observed upon repetitive stimulation, where VGLUT-pH exocytosis responses were diminished compared to wild-type (Fig 8F). This effect was temporary and recovered within ∼1 min.

## Discussion

Physiological sensory stimuli elicit different patterns of synaptic vesicle exocytosis and retrieval in the AWC^ON^ and ASH sensory neurons, as inferred from real-time changes in VGLUT-pH fluorescence. The ASH neuron has two distinct synaptic states: a basal state with low calcium levels and low exocytosis, and a stimulated state with increased exocytosis and high calcium levels. The AWC^ON^ neuron has three states: a basal state with tonic exocytosis and retrieval, an odor-evoked low-calcium state in which exocytosis is suppressed but retrieval continues, and a transient state after the removal of a long-duration odor stimulus with high calcium, accelerated exocytosis, and accelerated endocytosis.

Signaling at the *C. elegans* neuromuscular junction is dominated by graded synaptic release (Liu et al., 2009), consistent with the absence of sodium-based action potentials in nematode neurons (Goodman et al., 1998). A higher level of release can be evoked by direct electrical stimulation (Richmond et al., 1999), but mutant analysis suggests that evoked release is only weakly correlated with physiological function at the NMJ (Francis et al., 2005; Martin et al., 2011). In AWC^ON^, odor addition and subsequent odor removal represent physiological stimuli that are associated with tonic, suppressed, and evoked release. Interestingly, we saw no evidence of tonic synaptic release from ASH at rest, in contrast with the evidence for tonic release from AWC^ON^ and motor neurons.

### Cell type-specific regulation of the synaptic vesicle machinery in sensory neurons

The SNARE complex is required at all known synapses, but the detailed functions of the highly conserved SNARE regulators are still being determined. We found that the requirements for SNARE regulators differed between neurons. Both AWC^ON^ and ASH had a partial requirement for *unc-10/RIM.* However, *unc-13* was essential for all VGLUT-pH responses in ASH, but *unc-18* was not, whereas AWC^ON^ was more strongly dependent on *unc-18* relative to *unc-13.* Thus the synaptic requirement for *unc-13* and *unc-18* may differ across different synapses or conditions, even in cells in which both genes are expressed and active (Atwood and Karunanithi, 2002; Crawford and Kavalali, 2015; Kasai et al., 2012).

In *unc-13* mutants, AWC^ON^ did not respond to odor addition with a normal decrease in VGLUT-pH, but did exhibit some exocytosis immediately after odor removal. These dynamics suggest that tonic AWC^ON^ release cannot be maintained in *unc-13(lf)* mutants, but synaptic vesicles can be released after a large calcium influx. This result, among others, suggests that the underlying difference between ASH and AWC^ON^ cannot be explained entirely by the tonic-evoked distinction: in ASH, *unc-13* is required for evoked release, but in AWC^ON^, it is required for tonic synaptic exocytosis.

A striking difference between AWC^ON^ and ASH was their dependence on the protein kinase C epsilon homolog PKC-1. *pkc-1* mutants had severe defects in VGLUT-pH mobilization in AWC^ON^, resembling mutations in the SNARE complex. By contrast, exocytosis in ASH in *pkc-1* mutants was nearly normal, although a subtle defect could be uncovered by pulsing ASH with multiple stimuli.

Previous studies at the neuromuscular junction indicated that *pkc-1* affects neuropeptide release but not fast synaptic transmission from cholinergic and GABAergic motor neurons (Sieburth et al., 2007). Moreover, *pkc-1* mutants have near-normal locomotion, contrasting with the severe locomotion defects in SNARE mutants, providing further evidence of residual fast synaptic transmission at the neuromuscular junction. We suggest that the motor neurons, like ASH, can mobilize synaptic vesicles without *pkc-1,* in contrast with AWC^ON^ where *pkc-1* has a substantial role.

Protein kinase C is a well-established potentiator of neurotransmitter release at mammalian synapses (Wierda et al., 2007), a result consistent with those observed here. Interestingly, one of the targets of mammalian PKC is the homolog of UNC-18, Munc18-1, paralleling our observation that AWC^ON^ exocytosis has a strong requirement for both UNC-18 and PKC-1. In mouse hippocampal neurons, Munc18-1 clustering at synapses is regulated by activity, calcium influx, and protein kinase C phosphorylation, and correlates with synaptic strength (Cizsouw et al., 2014). At the mouse Calyx of Held, Munc18-1 PKC phosphorylation sites are important for post-tetanic potentiation, a form of plasticity that enhances neurotransmitter release (Genc et al., 2014). *C. elegans* UNC-18 shares consensus PKC phosphorylation sites, which may be phosphorylated by PKC-2, a different PKC, in thermosensory neurons (Edwards et al., 2012). While the link between PKC and UNC-18 is intriguing, PKC is likely to have multiple synaptic targets; for example, it phosphorylates the calcium sensor synaptotagmin-1 at hippocampal synapses to potentiate synaptic vesicle release (de Jong et al., 2016).

*unc-13* and *unc-18* mutants have severe locomotion defects and cannot be easily tested for sensory behaviors, but the more agile *pkc-1* mutants have been shown to be repelled, rather than attracted, by odors sensed by AWC^ON^ (Tsunozaki et al., 2008). Their responses to temperature sensed by AFD neurons can also have a reversed valence, with attraction to high temperatures that are normally repulsive (Luo et al., 2014; Okochi et al., 2005). The behavioral reversal in sensory responses could be related to dynamics of residual glutamate signaling, or it could result from an alternative form of neurotransmission such as neuropeptide release.

### Activity-dependent cytoplasmic pH decreases in glutamatergic neurons

Stimuli that evoke calcium increases in AWC^ON^ and ASH result in decreased cytoplasmic pH, as reported by pHluorin fluorescence changes. One possible source of the cytoplasmic change is acidosis evoked by increases in calcium levels, possibly due to proton mobilization during calcium extrusion by the plasma membrane calcium ATPase (PMCA) (Rossano et al., 2013; Schwiening and Willoughby, 2002; Trapp et al., 1996; Zhang et al., 2010). During periods of elevated calcium levels, pumps are active and acidify the neuron by exchanging protons for calcium ions; when the activity of the neuron decreases and calcium is extruded, activity of the pumps decreases, and the neuron returns to baseline pH levels (Trapp et al., 1996). Consistent with calcium driven acidosis, pH changes were faster in the axon than in the soma, like stimulus-induced calcium dynamics.

The stimulus-induced, exocytosis-independent cytoplasmic pH changes we observed in AWC^ON^ and ASH are worth future study, and may have unintended technical effects on other imaging experiments. For example, GCaMP fluorescence increases by as much as 10% for a 0.1 pH unit increase at physiological pH (Kneen et al., 1998; Nakai et al., 2001), and therefore stimulus protocols that alter cytoplasmic pH may confound GCaMP imaging in the same cells.

### Synaptic vesicle retrieval in glutamatergic neurons

In simple stimulus protocols, ASH neurons had a faster apparent rate of synaptic vesicle retrieval – the combination of endocytosis and re-acidification -- than AWC^ON^ neurons. However, the distribution of AWC^ON^ decay rates was shifted to faster timescales after a previous stimulation with odor (the acceleration effect), producing a distribution similar to that of ASH. The acceleration effect in AWC^ON^ endocytosis may be mediated by the large calcium influx generated by the removal of the first odor pulse, as calcium affects endocytosis in other species (Leitz and Kavalali, 2016; Neves et al., 2001; Sankaranarayanan and Ryan, 2001). Alternative or additional sources of this activity-dependent signal in AWC^ON^ include cGMP, which can accelerate endocytosis (Petrov et al., 2008) or cytoplasmic alkalinization (Zhang et al., 2010), as the AWC^ON^ axon becomes significantly alkalinized during odor stimulation.

The average basal and accelerated time constants of synaptic vesicle retrieval in AWC^ON^ are ∼18s and ∼8s, within the range reported in other systems. For example, the time constant of endocytosis in hippocampal neurons at physiological temperatures is ∼6s (Balaji and Ryan, 2007), and in goldfish retinal OFF-bipolar cells, the time constants for fast and slow endocytosis measured using a capacitance clamp were 1s and >10s, respectively (Neves and Lagnado, 1999). We did not detect signals on the timescale of the ultrafast endocytosis observed in *C. elegans* motor neurons and mammalian neurons (Watanabe et al., 2013a; Watanabe et al., 2013b). If ultrafast endocytosis is present in these sensory neurons, the time constant we measure is likely to represent subsequent re-acidification.

The AP180/pCALM homolog UNC-11 is required for efficient vesicle retrieval in AWC^ON^ and ASH. Our results are consistent with current models of AP180/CALM action that emphasize a role in clathrin-dependent sorting at a stage after endocytosis, but before the generation of mature, acidic synaptic vesicles (Soykan et al., 2017; Watanabe et al., 2014). It is worth noting that this protein probably has multiple functions in vesicle generation, as it affects both the molecular composition and size of synaptic vesicles (Koo et al., 2015; Koo et al., 2011; Nonet et al., 1999; Zhang et al., 1998).

### Conserved neuronal cell types in divergent animals

How similar, and how divergent, are neuronal cell types in different animals? In genetics, the concept of orthologous genes serves as a valuable framework for cross-species comparisons; whether such orthologous relationships apply to neuronal cells is a subject of debate (Vergara et al., 2017). The argument for cell type conservation has been mainly based on developmental transcription factors, rather than mature neuronal properties (Vergara et al., 2017). Here, we found that AWC^ON^ neurotransmission, like its sensory signaling, resembles that of vertebrate photoreceptor neurons. Both AWC^ON^ and photoreceptors have basal sensory activity that is suppressed with stimulation, cGMP-based transduction machinery, and circuitry that bifurcates into two streams of ON/OFF neurons (Chalasani et al., 2007). We observed that these similarities extend to the synaptic dynamics of AWC^ON^ and zebrafish and goldfish OFF-bipolar neurons, which resemble photoreceptors (Morgans, 2000; Odermatt et al., 2012). Both AWC^ON^ and OFF-bipolar neurons have fast and slow models of synaptic vesicle exocytosis and retrieval that are modulated by neuronal activity (Neves et al., 2001; Neves and Lagnado, 1999). Moreover, synaptic vesicle priming in photoreceptor and bipolar neurons of the mammalian visual system has been suggested to be largely independent of *Munc-13* (Cooper et al., 2012), and we observed residual synaptic vesicle release from AWC^ON^ in *unc-13(lf)* mutants. The extensive similarity between vertebrate retinal neurons and *C. elegans* olfactory neurons suggests that they are evolutionarily conserved cell types.

The differences between neuronal cell types across animals are also very substantial – for example, most *C. elegans* neurons do not have sodium-based action potentials (and neither do vertebrate photoreceptors). It remains to be seen how widely the idea of orthologous cell types holds, but it makes specific predictions. For example, we found that ASH and AWC^ON^ sensory neurons had different synaptic dynamics and molecular requirements: are ASH neurons similar to vertebrate nociceptors in their synaptic properties, as they are in their sensory use of TRPV channels? If neurons do fall into conserved classes, the convergence of genetics, behavior, and imaging tools such as pHluorins in *C. elegans* provide an avenue to uncovering their basic properties and underlying molecular mechanisms with single-cell resolution *in vivo*.

## Methods

### *C. elegans* culture

*C. elegans* strains were maintained under standard conditions on NGM plates at 21-22°C and fed OP50 bacteria (Brenner, 1974). Wild-type animals correspond to the Bristol strain N2. Transgenic lines were generated using standard methods by injecting young adult hermaphrodites with the desired transgene and a co-injection plasmid that expresses a fluorescent marker. In some cases, empty vector was included to increase the overall DNA concentration to a maximum of 100 ng/ul. A full strain list appears in Supplemental Methods.

### Fluorescence imaging of reporter proteins

GCaMP reporters were chosen to match the dynamic range of signaling in the relevant neuron and compartment. The higher-affinity GCaMP5A protein detects both calcium increases and decreases in AWC^ON^ cell body, whereas the lower-affinity GCaMP3 protein responds to the higher peak calcium levels in AWC^ON^ axons and the ASH cell body.

All imaging was conducted on a Zeiss Axiovert 100TV wide-field microscope. Images were acquired through a 100x 1.4NA Zeiss APOCHROMAT objective onto an Andor ixon+ DU-987 EMCCD camera using Metamorph 7.7.6-7.7.8 acquisition software. Imaging was conducted in custom-built microfluidic chambers designed for calcium imaging (Chronis et al., 2007). TIFF time-stacks were acquired at 5 frames per second (fps) using a 200 msec acquisition time. For AWC^ON^ GCaMP5 cell body recordings, TIFF time stacks were acquired using a 40x objective at 10 fps, 100 msec acquisition-time. Animals were age-synchronized by picking L4s onto fresh NGM OP50 seeded plates 12-18 hours before experiments. Imaging experiments were conducted in S basal buffer (Brenner, 1974). For recordings of AWC^ON^ activity, animals were starved for 20-30 minutes in an S basal bath prior to loading into the microfluidic chamber. For recordings of ASH activity, a 90-second recording was performed before any stimulation to allow animals to adapt to the blue light used for imaging. To further prevent animal movement during imaging experiments, animals were paralyzed with 1 mM (-)-Tetramisole hydrochloride (Sigma-Aldrich) during acquisition (Gordus et al., 2015). After loading in the microfluidic chambers, animals were allowed to acclimate for 5 minutes before imaging. Details of methods and data analysis appear in Supplemental Methods.

Butanone (Sigma) or NaCl (Fisher) stimuli were prepared fresh on the day of the experiment from pure stock solutions. The final butanone concentration was 11.2 uM (10^-6^ dilution, prepared by serial 10^-3^-fold dilutions) and the final NaCl concentration was 500 mM. Stimulus and control solutions were prepared in S basal buffer in amber glass vials. Control buffer and stimuli were delivered via reservoirs in 30 mL syringes (Fisher).

### Statistical analysis and curve fitting

All statistical tests are indicated in figure legends, and were performed using Prism 7 GraphPad software. Additional supporting statistical analysis for each figure can be found in the corresponding Source Data files. Fitting of decay curves was performed in Matlab using the ‘fit’ function with a custom ‘fitType’ equation. Traces that could not be fit due to high noise or did not exhibit a decay were removed from the analysis and is reflected in the change in n reported in the figure legends (see also Supplemental Methods). To compare the fit of different exponential models, we used the Akaike’s Information Criterion (AIC) as described in (Motulsky and Christopoulos, 2004) and Supplemental Methods.

### Molecular biology

For cell-specific expression in AWC^ON^ and ASH we used the promoters *str-2* and *sra-6,* respectively. The VGLUT-pH expression construct was created by subcloning super-ecliptic pHluorin into the first luminal loop domain of the *C. elegans* vesicular glutamate transporter *eat-4,* based on homology to mammalian VGLUT-1 (Voglmaier et al., 2006). Using site-directed mutagenesis (Stratagene quickchange protocol) we inserted a KPN-1 restriction site into the *eat-4.a* cDNA (wormbase CDS ZK512.6a) after the conserved glycine residue at position 106. Super-ecliptic pHluorin was inserted into this site using primers that added a 14 amino acid linker.

Forward primer:

*‘5-GAATCGTAGGTACCTCTACCTCTGGAGGATCTGGAGGAACCGGAGG ATCTATGGGAAGTAAAGGAGAAGAACTTT-3’*

Reverse primer:

*‘5-GAATCGTAGGTACCTCCGGTTCCTCCAGATCCTCCGGTTCCTCCGG TTCCTCCACCGGTTTTGTATAGTTCATCCA-3’*

syGCaMP was created by fusing GCaMP3 to the C-terminus of synaptogyrin-1 (*sng-1)*. *sng-1 cDNA* (wormbase CDS T08A9.3) was isolated from a N2 whole worm cDNA library and subcloned into a pSM expression vector containing GCaMP3 using AflII and SacII restriction enzymes. *sng-1* was fused to GCaMP3 through a 6x Glycine-Serine linker.

Forward primer: *5’- CAAATGATGACAGCGAAGTGGCTTAAGCATGGTATTGATAT CTGAGC-3’*

Reverse primer:*5’- GAATCGTAccgcggGAACCACTACCACTACCataaccatatcct tccgactga-3’*

To create CD4-pH, Super-ecliptic pHlourin was localized to the extracellular surface by fusion to a modified form of CD4 (Feinberg et al., 2008). CD4-pH was produced by exchanging the spGFP 1-10 in the pSM vector CD4-2::spGFP1-10 (Feinberg et al., 2008) for super-ecliptic pHluorin using the restriction sites Nhe1 and Sal1. The inserted super-ecliptic pHluorin contained the N-terminal linker domain: Gly-Gly--Gly--Gly--Gly-Ser-Gly--Gly--Gly--Gly-Ser.

AWC^ON^ *pkc-1* rescue: *pkc-1.a* cDNA (corresponding to wormbase’s CDS F57F5.5a sequence) was isolated from N2 whole worm cDNA libraries and subcloned into the pSM expression vector using Nhe1 and Kpn1 restriction sites. AWC^ON^ expression was achieved using the *str-2* promoter (vector str-2:pkc-1.a_cDNA:sl2:mCherry). Expression was confirmed for each animal tested checking for co-expression of mCherry. *pkc-1* cDNA isolation primers:

Forward:

*‘5-GAATCGTAGCTAGCATGCTGTTCACAGGCACCGTGC-3’*

Reverse:

*‘5-GAATCGTAGGTACCTTAGTAGGTAAAATGCGGATTGA-3’*

## Acknowledgements

We thank Aditya Rangan for guidance in kinetic modeling, Andrew Gordus for sharing his image analysis code, Tim Ryan and Jeremy Dittman for extensive discussions of synaptic imaging, and Johannes Larsch, Aylesse Sordillo and Sagi Levy for discussions and comments on the manuscript. This work was supported by the Howard Hughes Medical Institute.

---

**Supplemental Figure 1.**
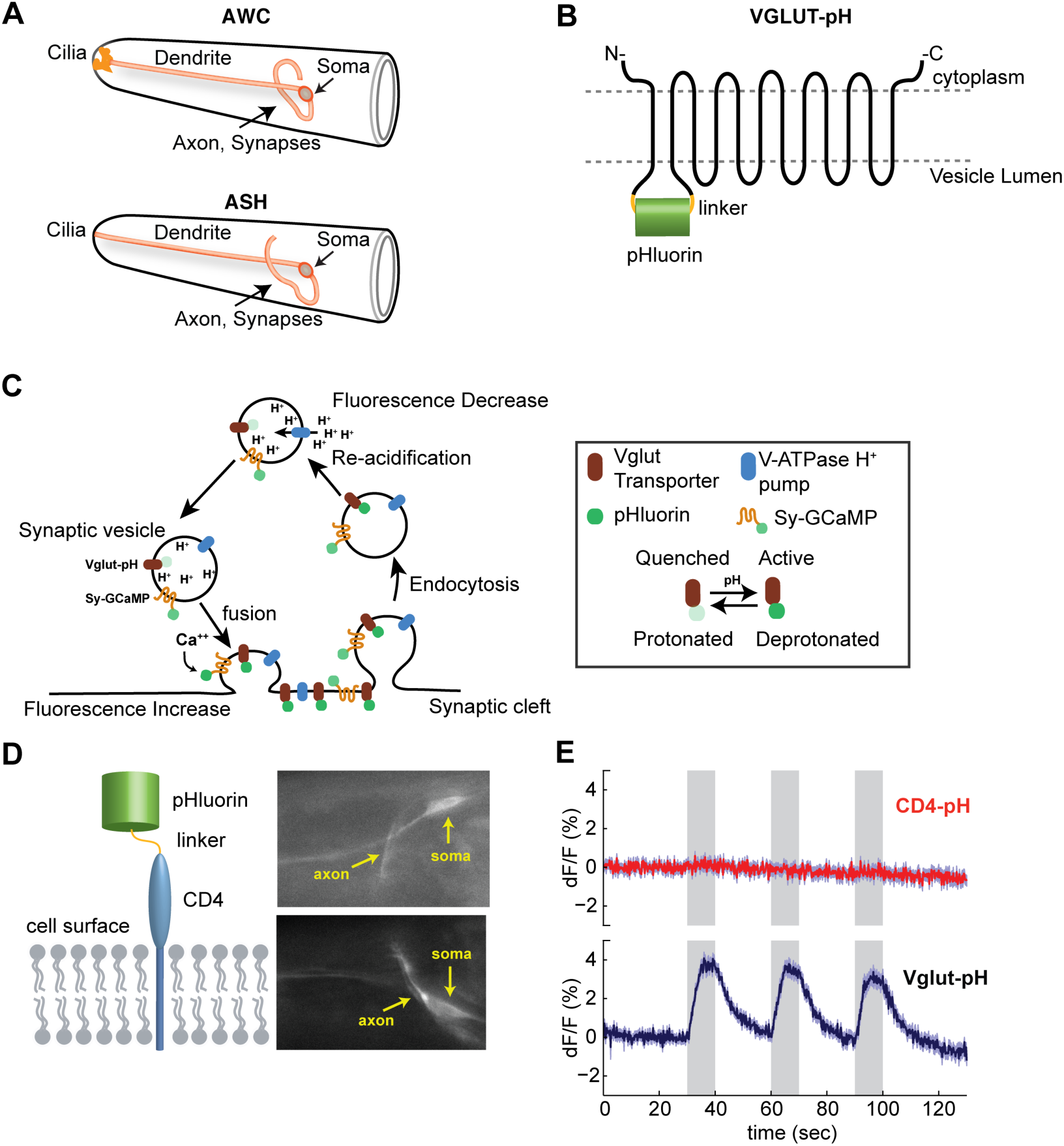
Anatomy of AWC^ON^ and ASH and illustration of synaptic reporters. **(A)** Anatomical diagram of AWC and ASH neurons. Redrawn from www.wormatlas.com. **(B**) Diagram of VGLUT-pH reporter, with pH-sensitive fluorophore inserted into a lumenal domain of the protein. **(C)** Diagram of simplified synaptic vesicle cycle, incorporating synaptic activity reporters used here. VGLUT-pH reports pH changes of the synaptic vesicle lumen associated with exocytosis. syGCaMP, a fusion to the synaptic vesicle protein synaptogyrin, reports changes in calcium concentration at the cytosolic face of synaptic vesicles. These reagents are used separately. **(D)** LEFT: Diagram of CD4-pH construct used to target pHluorin to the extracellular surface of ASH. RIGHT: Two animals expressing CD4-pH in ASH under the *sra-6* promoter. ASH axon and cell body (soma) indicated. Images are from recordings in (E) and are averages of the first 5 frames. **(E)** TOP: Average response of ASH CD4-pH (n=33 trials, 11 animals, 3 trials each). No response was detected. BOTTOM: Average response of ASH VGLUT-pH control (n=39 trials, 13 animals, 3 trials each). Gray bars mark stimulus periods (500 mM NaCl). Shading = S.E.M.

**Supplemental Figure 2.**
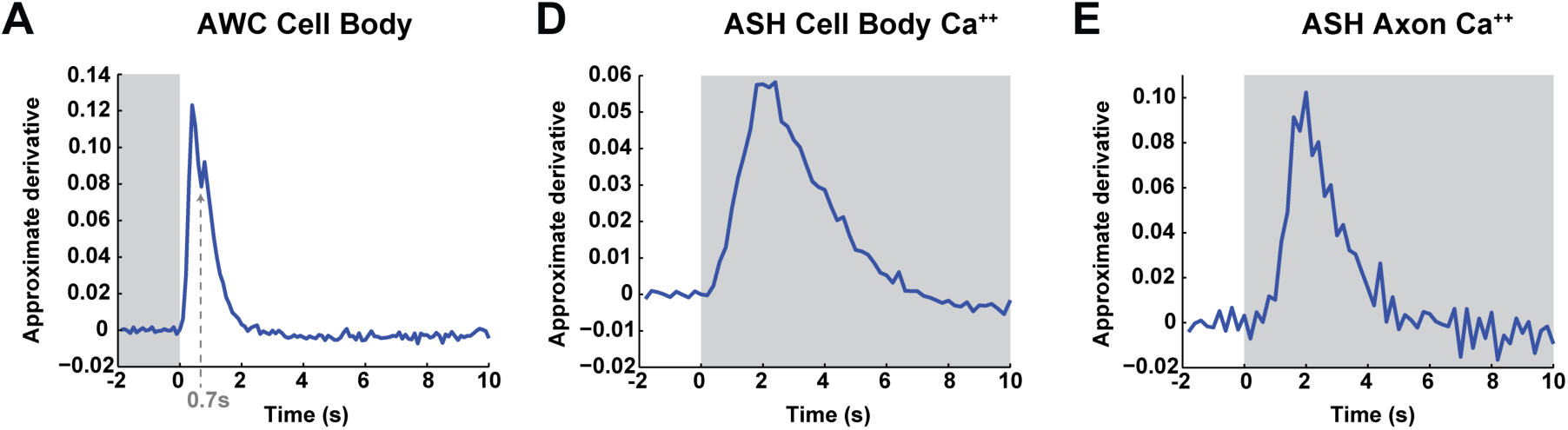
**(A)** Time derivative of the mean AWC^ON^ GCaMP5 signal in the cell body after butanone removal (60s odor stimulus, 11.2uM). A transition between two positive influx rates may occur ∼0.7s after odor removal, similar to the transition observed in synaptic calcium by syGCaMP recordings in Figure 2G. Average from 47 trials from 16 wild-type animals. **(B,C)** Time derivative of the mean ASH GCaMP3 signal at the cell body **(B)** and axon **(C)** during a 500 mM NaCl stimulus. No heterogeneity of influx rates was detectable with this sensor. Data from Fig 1B.

**Supplemental Figure 3.**
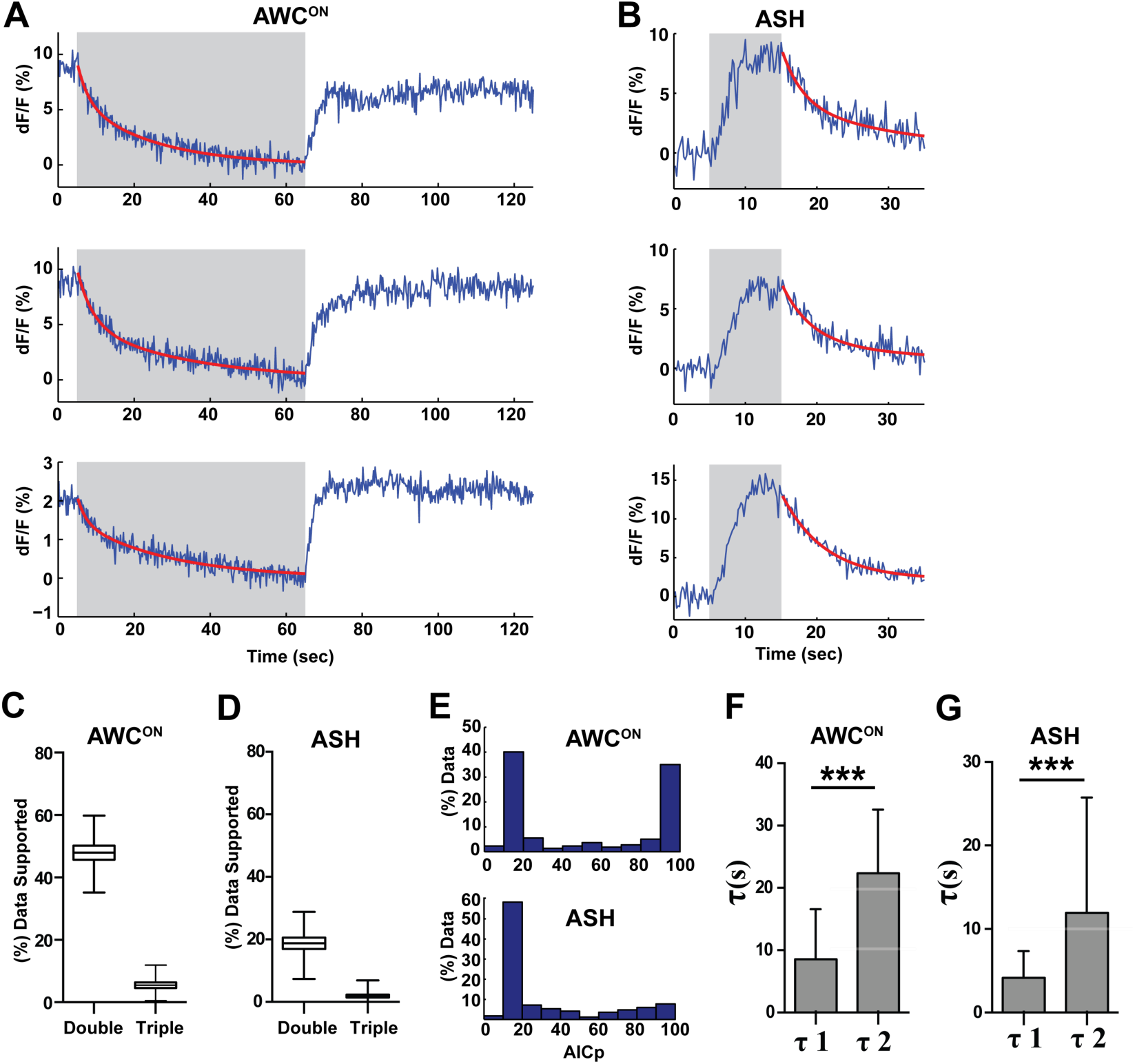
A proportion of VGLUT-pH recordings are consistent with multiple retrieval time constants. **(A,B**) Three individual examples of AWC^ON^ (A) or ASH (B) VGLUT-pH decays that fit a double exponential decay model with greater than 95% likelihood based on AICp (see Methods). **(C,D)** Percent of VGLUT-pH data that fits a single versus double, or double versus triple exponential model with greater than 50% likelihood (AICp > 50%). Box and whisker plot created by 10,000 sampled bootstrap sampling. **(E)** Frequency histogram of AICp-based likelihood values for double exponential fits to VGLUT-pH data. An AICp value of >50 supports a double exponential fit with increasing likelihood; an AICp of less than 50 supports a single exponential with increasing likelihood. (C-E) AWC^ON^ data: n=218 (76 animals, 2-3 trials each). ASH data: n=168 (56 animals, 2-3 trials each). **(F,G)** Average decay constants from double exponential fits. Note similarity of two AWC^ON^ timescales to effects of plasticity in Fig 7F. Fits that minimized tau1 or tau2 to ∼zero were excluded from the average. AWC^ON^ data: n = 211 (76 animals, 2-3 trials each). ASH data: n = 125 (56 animals, 2-3 trials each). Error bars = standard deviation. t-test p <0.0001. ASH and AWC^ON^ data from (Fig 1E,G). AWC^ON^ stimulus = 11.2 uM butanone. ASH stimulus = 500 mM NaCl.

**Supplemental Figure 4.**
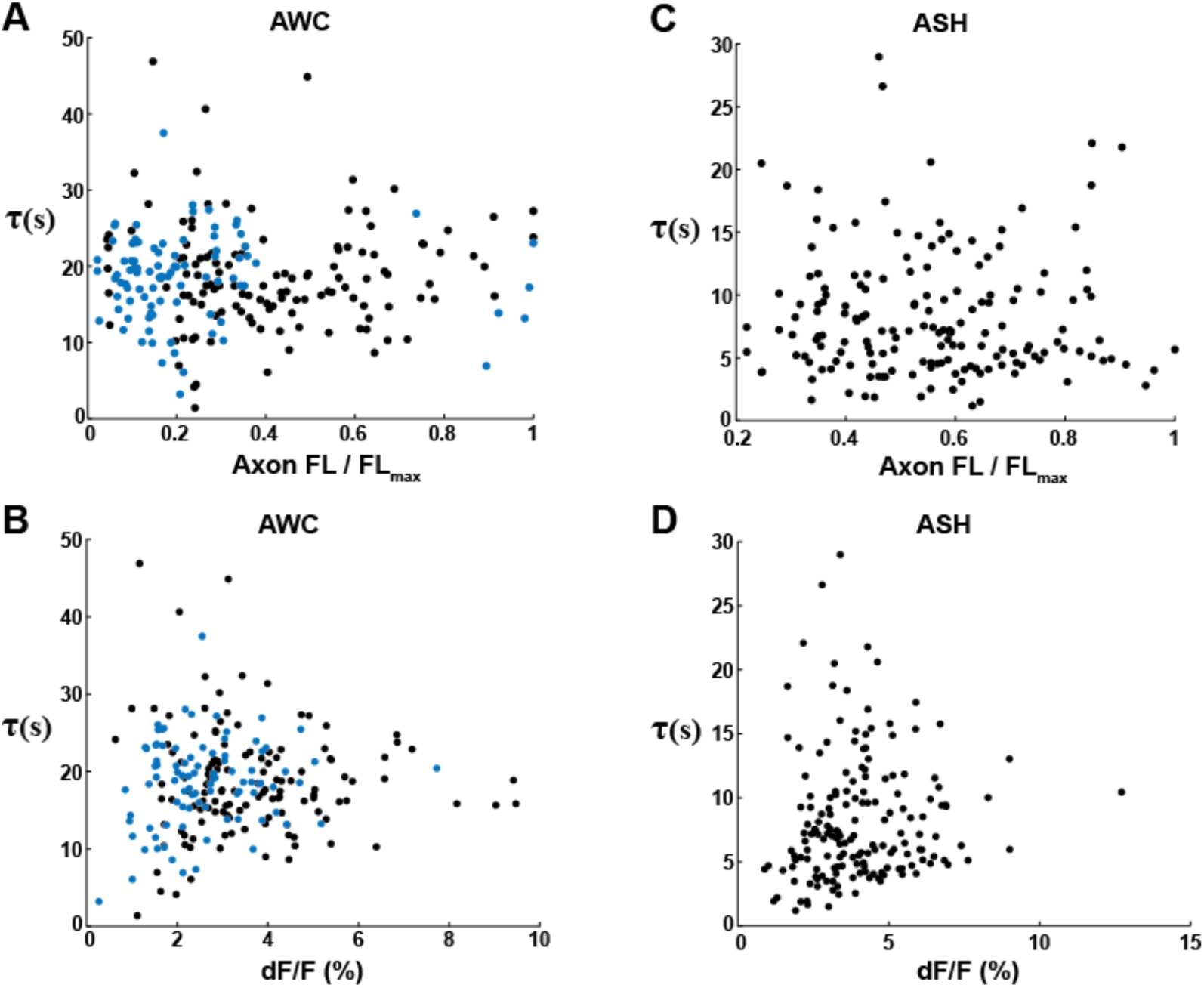
VGLUT-pH decay time constants are not correlated with expression level or response magnitude. **(A)** Scatter plot of AWC^ON^ VGLUT-pH decay time constants versus baseline axon fluorescence intensity. Two different AWC VGLUT-pH transgenic lines with different expression levels are represented by the black and blue points. Axon intensities were normalized to the brightest axon detected in the line (FL_MAX_). Time constants are from single exponential fits, as in Figure 6. Correlation coefficient =0.01, p=0.89, not significant. **(B)** AWC^ON^ decay time constants plotted against the magnitude of basal release, dF/F_0_ prior to stimulus onset; F_0_ is fluorescence level prior to stimulus removal. Correlation coefficient =0.024, p=0.73, not significant. **(C)** Scatter plots of ASH VGLUT-pH decay time constants versus baseline axon fluorescence intensity. Correlation coefficient = −0.029, p =0.72, not significant. **(D)** ASH decay time constants after stimulus removal, compared to peak dF/F during stimulus. Correlation coefficient =0.058, p =0.46, not significant. AWC^ON^ n=218 (76 animals, 2-3 trials each). ASH n=168 (56 animals, 2-3 trials each).

**Supplemental Figure 5.**
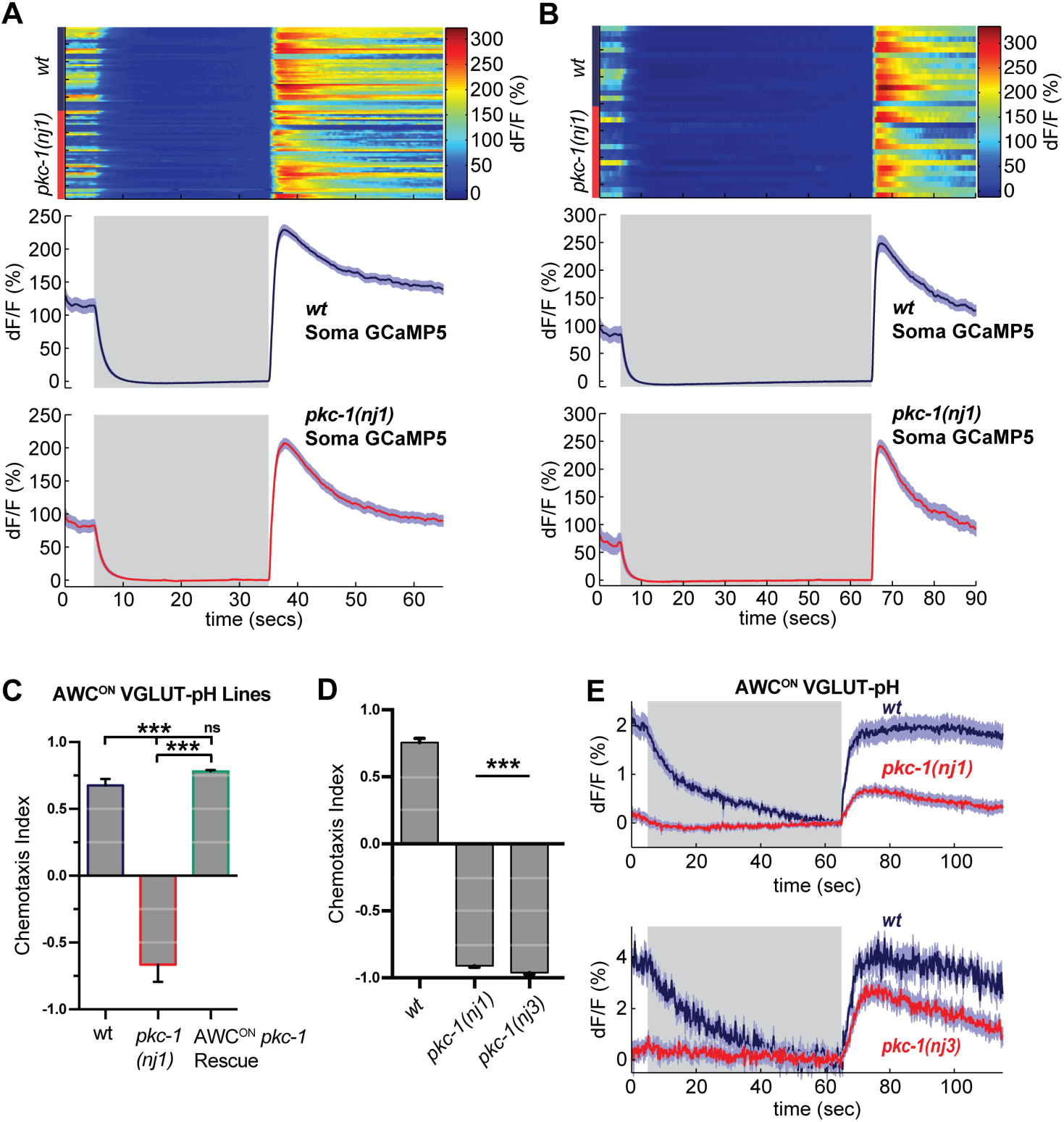
*pkc-1(lf)* alters synaptic release downstream of calcium influx. **(A,B)** AWC^ON^ GCaMP5 responses to 30s (A) and 60s (B) pulses of butanone, measured at the soma. Top panels: Heat map of individual trials. Bottom: average across trials. (A) *pkc-1(nj1)* n = 50 (17 animals, 2-3 trials each), *wt* n = 47 (16 animals, 2-3 trials each). (B) *pkc-1(nj1)* n = 16 (16 animals), *wt* n = 16 (16 animals), 1 trial each. **(C,D)** Butanone chemotaxis index (See Supplemental Methods) of AWC^ON^ VGLUT-pH transgenics used in Fig 8D, showing that expression of *pkc-1* cDNA in AWC^ON^ under the *str-2* promoter rescues *pkc-1(nj1)* chemotaxis. n = 3 population assays per genotype. One-way ANOVA, *** p <0.0001. ns = not significant. (D) Butanone chemotaxis of *pkc-1(nj1)* and *pkc-1(nj3)* mutants. *pkc-1(nj1)* n = 26, *pkc-1(nj3)* n = 6, *wt* n = 29 population assays. One-way ANOVA, *** p <0.0001 compared to wt. ns= not significant. Error bars = S.E.M. **(E)** VGLUT-pH responses in *pkc-1(nj1)* and *pkc-1(nj3)* mutants. *pkc-1(nj1)* data is from Fig 8B. Bottom panel: *pkc-1(nj3)* n = 18 (6 animals, 3 trials each), *wt* n = 6 (2 animals, 3 trials each). Gray bar = stimulus, 11.2 uM butanone. Shading = S.E.M.

**Supplemental Figure 6.**
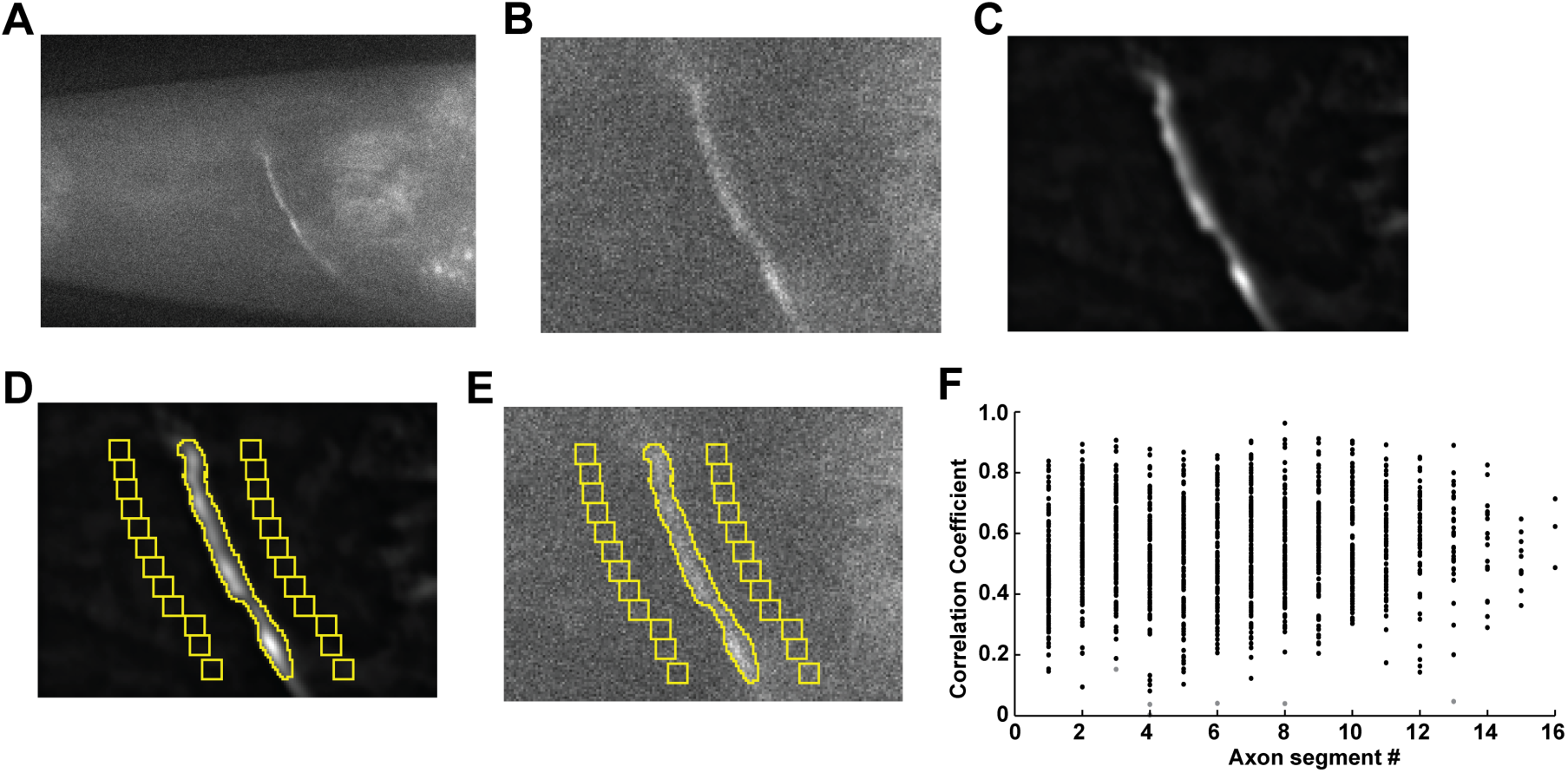
VGLUT-pH data acquisition. **(A)** Single plane of AWC^ON^ VGLUT-pH recording after drift-registration correction. **(B)** Zoomed view around the axon in (A). **(C)** Mean intensity projection after the recording was processed by rolling background subtraction and 3D-averaging. **(D)**Selection of axon and background ROIs. **(E)** Intensity measurements are then taken from the raw drift-corrected recording. **(F)** ROI segments along the axon correlated with overall mean activity of the axon. Each point represents the correlation coefficient of an axon segment with the mean signal of the axon during a standard recording (see text). Data from 120 AWC^ON^ VGLUT-pH axons stimulated with 11.2 uM butanone. Each axon had 8 – 16 ROI segments. p < 0.000001 for all correlation coefficients except the points labeled in light gray, which were not significant p > 0.05.

## Supplemental Methods

## Additional information on imaging activity-dependent fluorescence reporters

All imaging was conducted on a Zeiss Axiovert 100TV wide-field microscope. Images were acquired through a 100x 1.4NA Zeiss APOCHROMAT objective onto an Andor ixon+ DU-987 EMCCD camera. Camera settings: 14-bit EM-GAIN enabled digitizer (3MHz); baseline clamped; overlapped recording mode; 0.3 uS vertical clock speed; binning = 1.

Images were cropped to fit around the head of animal. Most experiments used a pre-amplifier gain of 5x. Illumination was provided by a Lumencore SOLA-LE solid-state LED lamp. Illumination input was passed through a 1.3 ND filter. Narrow bandwidth blue light illumination (484-492 nm) was produced using the CHROMA 49904-ET Laser Bandpass filter set.

Stimulus triggering was performed through Metamorph via digital input from a National Instruments NI-DAQmx box to an Auotmate Valvebank 8 II actuator that triggered Lee Corporation solenoid valves. Custom journals specified pre-programmed recording parameters and performed automated file naming and storage. Most stimulation protocols involved multiple trials per animal, as detailed in figure legends. The interval length for all trials was 30 seconds in addition to the time recorded before or after the stimulation. For AWC^ON^ recordings in which the pulse length was varied, stimulation was ordered sequentially as follows: 6 trials with 10s pulses, 3 trials with 60s pulses, and a single 3 minute pulse. Order of stimulation did not appear to affect results.

## Quantification of fluorescence changes

dF/F was calculated as:

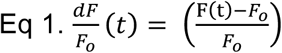

Where F (t) = is the fluorescence of a ROI minus the background at time (t) and F_o_ = fluorescence of the ROI minus the background at some reference time point or value.

For AWC^ON^, F_o_ was set to the mean background-corrected signal two seconds before odor removal. For ASH recordings, F_o_ was set to the mean background-corrected signal two seconds before the first stimulus for each trial. In each case F_o_ corresponded to a stable baseline within and across recordings.

## Acquisition of Fluorescence Measurements

To extract fluorescence measurements from VGLUT-pHluorin images, we developed custom semi-automated tracking software in ImageJ. Images were first corrected for x-y drift using image registration (Tseng et al., 2011), which places the axon at a specific set of image coordinates for the entire recording by shifting each frame in x and/or y. The microfluidic device prevents most z-plane drift, but images in which significant z-plane drift was detected were discarded. To aid in axon selection, images underwent rolling-ball background subtraction and then were averaged over space (2 pixels in x and y) and time (average of time-stack). The entire segment of the axon that was in view was then specified by the user and outlined by hand with the aid of pixel intensity thresholding. From this axon outline, intensity and pixel information was extracted from the raw drift-corrected recording along with local background measurements along the axon (Fig S6A-E). This process was also used to acquire measurements of AWC^ON^ syGCaMP images. For ASH, the *sra-6p* promoter is also expressed in the neurons ASI and PVQ, the axons of which are posterior to the ASH axon. VGLUT-pH fluorescence could also be detected in these other axons (mainly PVQ) but did not significantly respond to stimulation. In any given experiment, we recorded VGLUT-pH signals from a single ASH axon, either the on right or left side of the animal. Because of ASI and PVQ VGLUT-pH fluorescence, only the anterior background ROIs were used in analysis of ASH VGLUT-pH recordings.

For cell body measurements of GCaMP responses in AWC^ON^ and ASH we used a custom written ImageJ script written by Andrew Gordus (Gordus et al., 2015) to track cell body position and extract intensity measurements.

To validate using the whole axon segment as a single integrated measurement in VGLUT-pH experiments we measured VGLUT-pH responses from small equally spaced ROIs along the axon. These were obtained by taking the outlined axon segment as in Suppl. Fig 6E and cutting the axon into smaller ROI segments, each 8 pixels long in the y-axis, and performed a correlation analysis on a large dataset of AWC^ON^ VGLUT-pH recordings from wild-type animals that were stimulated with a 60 second pulse of 11.2 uM butanone. For 120 stimulated axons, all ROIs along the axon were positively correlated with the mean integrated axon fluorescence (Fig S6F). This correlation appeared to hold for all genotypes and conditions tested. There was substantial scatter in the strength of this correlation across different ROIs, potentially consistent with heterogeneity among synaptic regions (Fig S6F), but this possibility was not examined further.

## Background and bleaching correction of VGLUT-pHluorin signals

VGLUT-pH has a low baseline signal that is relatively close in intensity to the background autofluorescence generated by the surrounding tissue of the head. This became problematic when attempting to apply background subtraction and perform comparisons of fluorescence change. Most commonly, this is done with the deltaF function (dF/F) described above (Eq.1). The deltaF function normalizes the fluorescence change to its baseline intensity, allowing a comparison of signal changes between conditions with different absolute values. However, the deltaF function is highly sensitive to fluctuations in F_o_ when F_o_ is small. For VGLUT-pH, a further confound is created by reporter localization to parts of secretory pathway(s) that do not participate in synaptic release. These can be seen as small puncta that do not contribute to VGLUT-pH responses and, in rare events, these puncta can be observed undergoing translocation within the axon towards the cell body. These contribute to F_o_ but not to fluorescence changes. To prevent background correction from shifting F_o_ into a realm where small variations in F_o_ generate large fluctuations in deltaF, we subtract only the fluctuations in the background signal.

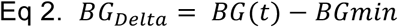

Where *BG =* the background at time (t) and *BGmin =* the minimum background value during the recording.

VGLUT-pH has very slow bleaching kinetics (Ariel and Ryan, 2010; Balaji and Ryan, 2007), essentially showing little to no bleaching over the timecourse of our recordings. This is likely the result of the reporter existing mainly within synaptic vesicles in the quenched state, and the rapid cycling of the fluorescent form back into this state. Unlike the VGLUT-pH reporter, background autofluorescence does show significant bleaching, and therefore, using this background signal for correction can result in an artificial increase in the VGLUT-pHuorin signal over time. To avoid this artifact, we corrected for background bleaching before performing background correction. Bleaching was assumed to be approximately linear and was modeled by line fitting using the Matlab function ‘polyfit.’ A threshold for specifying significant bleaching was set to a 0.5% drop in mean fluorescence intensity over 2 minutes of recording time. Bleaching was corrected by subtracting the linear fit from the background signal.

## Statistical analysis and curve fitting

All fitting of decay curves was performed in Matlab using the ‘fit’ function with a custom ‘fitType’ equation. For single exponential fits fitType = a1*exp (-x/tau1) + C. For double exponential fits fitType = a1*exp (-x/tau1) + a2*exp (-x/tau2) + C. For triple exponential fits fitType = a1*exp (-x/tau1) + a2*exp (-x/tau2) + a3*exp (-x/tau3) + C. In each case, all ‘a’ and ‘tau’ variables are fitted parameters and are constrained to be > 0. C = the baseline to which traces decayed, normalized to zero. Unless stated otherwise, fitting was applied to the entire decay period of the trace. For AWC^ON^ this corresponds to the odor-addition phase. For ASH, this corresponds to the 20 seconds immediately following stimulus removal. For comparison of AWC^ON^ decays with different stimulus durations (i.e. 60 second vs 20 second), we fit both traces using the initial 20 seconds to keep comparisons consistent. Decay constant averages reported for a given condition were performed as follows: each individual trace for a given data set was fitted individually, and each fit was plotted on its trace and then inspected. Traces that could not be fit due to high noise or did not exhibit decay were removed from the analysis. The decay constants for each condition were then collected and exported to Prism 7 for statistical analysis.

## Comparing exponential fits using AIC_C_

To compare models we used the Akaike’s Information Criterion (AIC) as described in (Motulsky and Christopoulos, 2004): 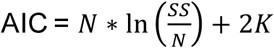, where N = number of data points, K = number of parameters, and SS= is the sum of the square of the vertical distances of the points from the curve. We used the corrected version for small 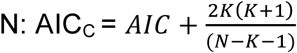 was performed for the fit on each individual trace. To determine of the relative likelihood of two models, we computed the probability that one model is more likely than the other 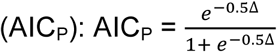, where Δ= the difference in AIC_C_ scores. All AIC based analysis was conducted in Matlab using custom scripts.

## *C. elegans* strains

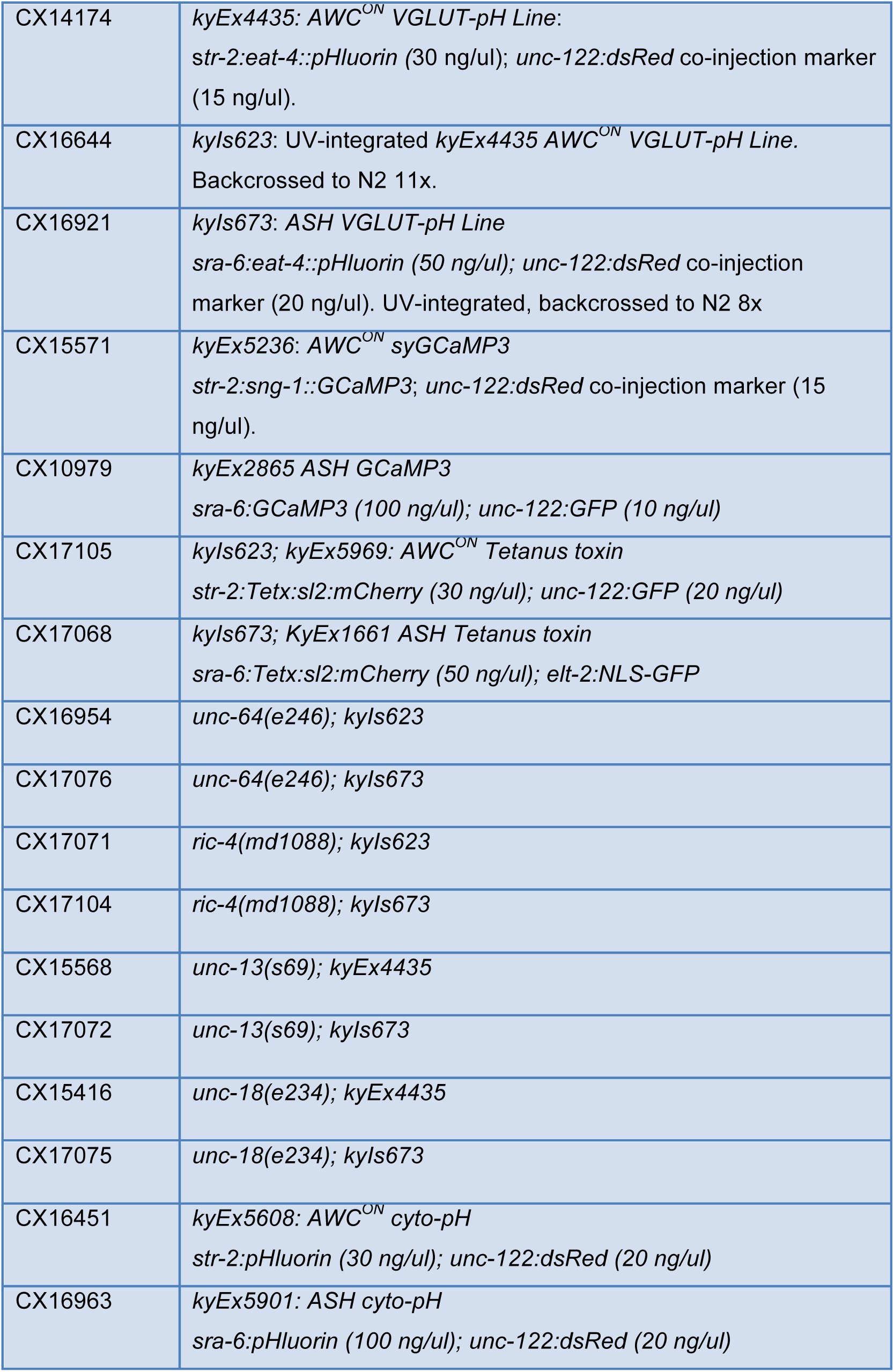
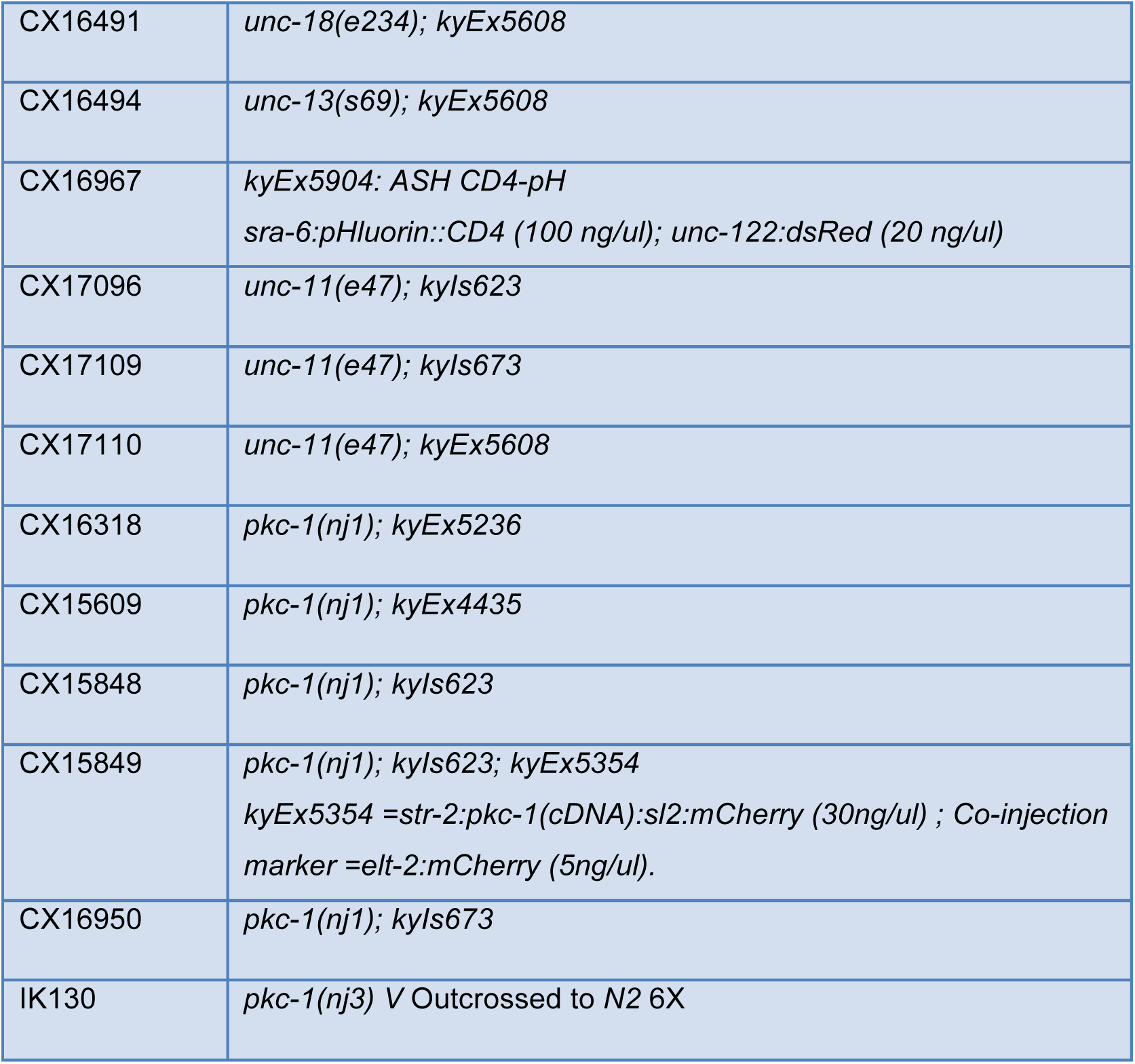

## Molecular identify of mutants

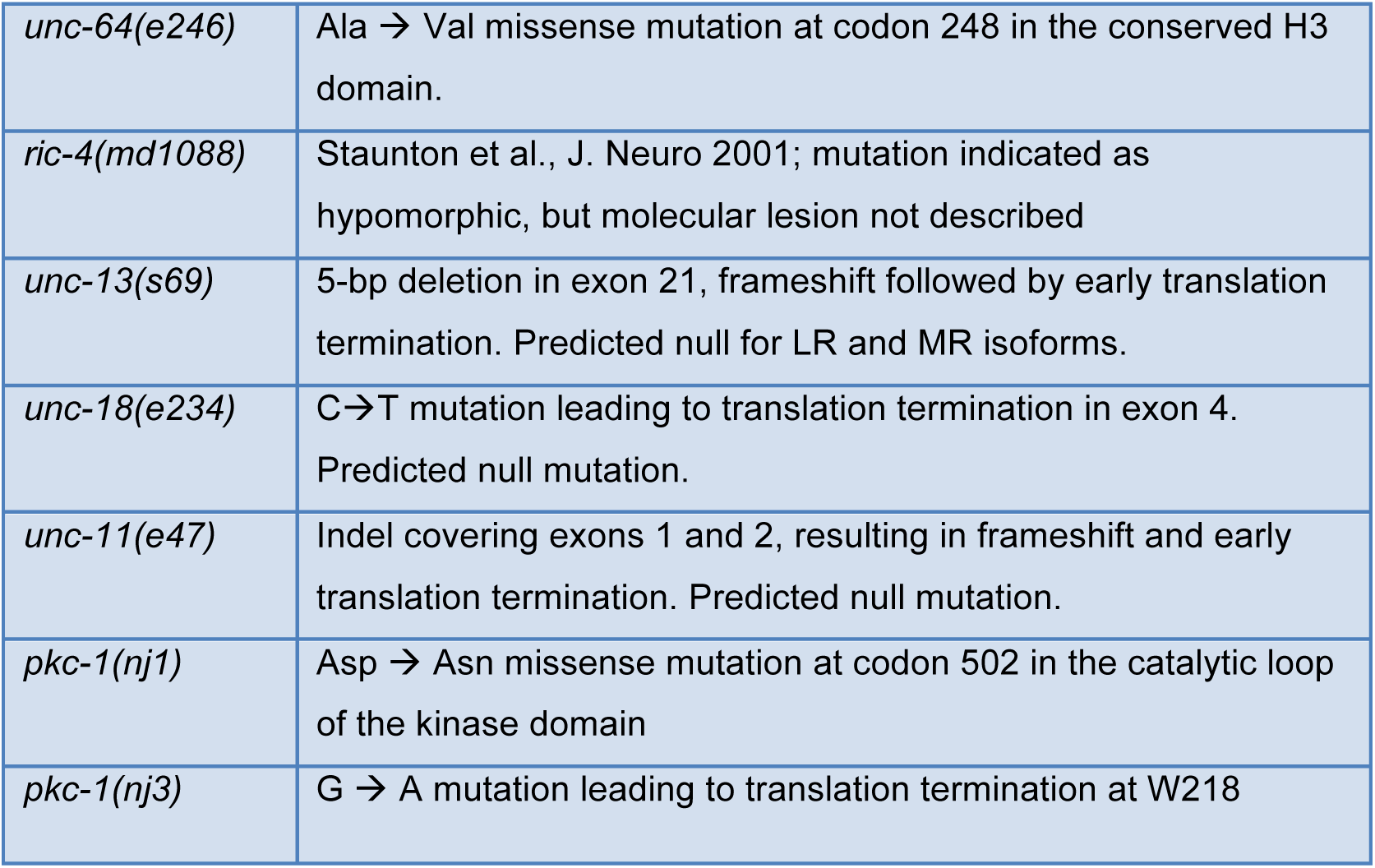

